# New peptides with immunomodulatory activity in macrophages and antibacterial activity against multiresistant *Staphylococcus aureus*

**DOI:** 10.1101/838201

**Authors:** Laura Andrea Barrero-Guevara, Natalia Bolaños, Miguel Parra, John Mario González, Helena Groot, Carolina Muñoz-Camargo

## Abstract

*Staphylococcus aureus* infections are a common concern world-wide due to the increasing number of bacterial strains with multiresistant properties to existing antibiotics, incrementing the need for novel molecules and therapy approaches for their treatment. This study evaluated the antibacterial and immunomodulatory activity of eight new peptides (AA, KS, NS, RN, AT, GF, KV and LK) as the basis for the search of new antibacterial and therapeutic agents for topic prevention and treatment against *S. aureus* infections. Here, there are characterized *in silico* eight new antimicrobial peptides. Their antibacterial activity against *S. aureus* and cytotoxic activity in mammalian cell lines were evaluated *in vitro* with the peptides individually and combined. Three of the peptides (GF, AT and AA) immunomodulatory activity was assessed in macrophages and under three scenarios: non-stimulation, *Escherichia coli* LPS stimulation and *S. aureus* lysate stimulation. Results showed that three peptides individually showed the best antibacterial activity against the *S. aureus* bacteria evaluated. The peptides presented immunomodulatory activity in THP-1 macrophages by displaying different profiles, increasing or decreasing four cytokines (IL-1β, TNF-*α*, IL-8 and CCL2 (MCP1)). This activity depended on the peptide concentration and the stimulation in which the macrophages were exposed to. Taken together, these results demonstrate the potential of these peptides to be used in further studies as novel antimicrobial molecules for the prevention and treatment of *S. aureus* infections.

## Introduction

Antimicrobial peptides are commonly defined as small, cationic molecules that consist of 10 to 50 amino acid residues found in a broad range of organisms (Zhang, 2016). The presence of antimicrobial peptides in the skin secretions of frogs has been confirmed decades ago. Such peptides are now considered part of the immune response of these and other animals (Conlon, 2011). Still, their precise role in the biology and survival of the organisms remains largely unknown, although their antimicrobial activity against bacteria (Groot *et al*. 2012), fungi (Rollins-Smith *et al*., 2002) and viruses (Muñoz *et al*., 2016) have long been studied. Recently, the description of peptides with (Yeung, Gellatly Hancock, 2011) immunomodulatory (Conlon *et al*., 2014 y Pantic *et al*., 2014), chemoattractant (Guaní-Guerra *et al*., 2010) and cytotoxic (Guaní-Guerra *et al*., 2010) activity, which has lead to rename them as host defense peptides.

In a worldwide panorama of increasing numbers of infections caused by multiresistant bacteria, it is important to keep searching for new therapeutic molecules (WHO, 2014). Taking into account the potential of the peptides secreted from frog’s skin glands as antibacterial and immunomodulatory molecules, they could be considered a promising alternative in clinical applications (Conlon *et al*. 2014). One of the purposes of immunomodulatory therapies is the control of infectious and non-infectious diseases. In this, coadjuvant molecules, such as antimicrobial peptides and pathogen-associated molecular patterns (PAMPs), regulate the immune response (Hancock *et al*, 2012).

*S. aureus* is one of the microorganisms which has caught the greatest attention worldwide, not only because of its role in nosocomial infections but also in community-acquired infections. These Gram-positive bacteria are responsible for many diseases ranging from simple skin infections to more serious conditions such as sepsis (Fournier Philpott, 2005). *S. aureus* can colonize the host as a commensal microorganism, however, it can also arise as a pathogen. Although *S. aureus* can infect many other tissues, the skin is where the majority of the infections develop (Kluytmans, Van Belkum Verbrugh, 1997). For the prevention and treatment of minor and secondary infections in soft tissues, mupirocin and bacitracin are the recommended topic agents (Bamberger Boyd, 2005). Nevertheless, recent studies have shown, that the resistance to these antibiotics is increasing worldwide (Patel, Gorwitz Jernigan, 2009). Taking into account the prevalence of *S. aureus* in human populations, the need to increase effort in the search of preventing measures, including topical options, becomes evident.

Studies have indicated, that the pathogenicity of *S. aureus* is caused by a great repertoire of toxins, exo-enzymes, adhesins, and proteins capable of immunomodulation (Fournier Philpott, 2005). Additionally, during the innate immune response against *S. aureus*, it has been found, that the severity of the infection varies according to the host’s differential secretion of cytokines (Brown *et al*., 2014). Specifically, in the skin, inflammation due to cytokines secreted by local macrophages exacerbates the severity of the bacterial infection (Montgomery *et al*., 2013 y Myles *et al*., 2013). Therefore, it has been proposed recently, that the *S. aureus* infection prevention should include immunomodulatory therapy (Wardenburg, Williams Missiakas, 2006 y Thangamani, Younis Seleem, 2015).

Hence, we aimed to evaluate the potential antibacterial and immunomodulatory activity of eight new peptides as the basis for the search of new antibacterial agents for the topic prevention and treatment against *S. aureus* infections. The general peptide characteristics, stability related features, sequence similarities with previously described proteins and antimicrobial peptides, as well as the antigenicity were analyzed *in silico* with a broad repertoire of bioinformatic tools. Then, the antibacterial effect of the peptides against *S. aureus* was identified *in vitro* and compared with that of commercialized antibiotics. Also, the cytotoxic effect of the peptides was evaluated in VERO fibroblasts, human monocytes (THP-1 cell line) and differentiated human macrophages (from THP-1 cell line). With these, the immunomodulatory effect of the peptides in differentiated human macrophages, stimulated with or without *E. coli* lipopolysaccharides (LPS) or *S. aureus* lysate, was identified by evaluating their capacity to enhance and/or inhibit the production of 13 different cytokines.

## Results

### General potential antimicrobial features of the eight new peptides showed important features which predict a promising antimicrobial potential

The positive net charge and the amphipathic composition are considered predictors of the peptides’ potential for presenting antimicrobial activity, as they might indicate their capacity to interact with living cell membranes. Although the eight peptides (AA, KS, NS, RN, AT, GF, KV and LK, named after their first and last amino acid of their sequence) used in this study had these characteristics, NS and AT peptides showed the most promising values (Table 1). Furthermore, according to the hydrophobic ratio, the peptides were classified as high hydrophobic peptides (AT, GF, KV and LK) and low hydrophobic peptides (AA, KS, NS and RN). Meanwhile, other features ranked the potential of the peptides differently, for example, the W-W Hydrophobicity indicated that the GF peptide might have a higher possibility to bind to a membrane, while NS might not stay attached (Table 1). Also, the Boman Index showed, that all the peptides evaluated in this study have a low potential to bind to other proteins. Finally, the Antimicrobial Peptides Database (APD) predicted that some of these peptides might form *α* helices or have sequences rich in rare amino acids, special features which some described antimicrobial peptides present (Table 1). In conclusion, although the eight peptides described in this study do not share the same characteristics, all predictions indicated their potential to be antimicrobial peptides.

**Table 1.** General potential antimicrobial features of the eight new peptides. *In silico* analysis using the Antimicrobial Peptides Database were conducted to identify general characteristics of antimicrobial peptides of the eight new peptides described in this study. All the peptides have consistently shown general characteristics of typical antimicrobial peptides, such as positive net charge and amphipathic composition. Nevertheless, the peptides were classified into two groups: high hydrophobicity peptides (AA, KS, NS and RN) and low hydrophobicity peptides (AT, GF, KV and LK). Furthermore, GF was identified as the peptide with the highest possibility to bind to membranes, while none of the peptides were considered to have the potential for binding to other proteins.

| ID | Isoelectric Point<sup>1</sup> | Net Charge<sup>2</sup> | Hydrophobic ratio<sup>2</sup> | GRAVY<sup>1,2</sup> | W-W Hydrophobicity (kcal/mol)<sup>2</sup> | Boman index (kcal/mol)<sup>2</sup> | Predicted Antimicrobial Features<sup>2</sup> |
| --- | --- | --- | --- | --- | --- | --- | --- |
| AA | 10 | +2 | 45 % | 0,36 | 2,54 | 0,07 | α-helix |
| KS | 10,3 | +3 | 25 % | -1,30 | 2,49 | 2,06 |  |
| NS | 10,18 | +4 | 35 % | -0,84 | 7,27 | 2,14 | α-helix |
| RN | 11 | +2 | 47 % | 0,73 | 0,92 | -0,26 | α-helix |
| AT | 10,3 | +3 | 53 % | 0,37 | 1,87 | 0,28 | α-helix |
| GF | 11 | +2 | 54 % | 0,89 | -0,94 | -0,43 | Rich in Prolines |
| KV | 8,75 | +1 | 50 % | 0,92 | 1,45 | -0,91 | α-helix |
| LK | 8,75 | +1 | 52 % | 1,06 | 0,78 | -1,22 | α-helix |

ID = Code given to each peptide according to the first and last amino acid of its sequence. Hydrophobic ratio= Total Hydrophobic Ratio as defined by APD. GRAVY= Grand Average Hydrophathy Value of the peptide. W-W Hydrophobicity= Wimley-White Whole-Residue Hydrophobicity of the peptide, calculated as the sum of whole-residue free energy of transfer of the peptide from water to lipid bilayer interface. Boman Index= Protein Binding Potential to other proteins. Predicted Antimicrobial Feature= APD Predictions of Antimicrobial Features. 1. ExPASy, Bioinformatics Resource Portal, ProtParam Tool (Gasteiger et al., 2005). 2. APD, APD3: Antimicrobial Peptide Calculator and Predictor (Wang & Wang, 2016)

### Predicted stability features of the new peptides *in vivo, in vitro*, in test tubes, in an intestine-like environment and in presence of proteolytic enzymes and chemicals indicate that the peptides are stable in proteolytic environments, but susceptible to peptidases originating from bacteria of clinical importance

The half-life predictions obtained according to the N-terminal amino acid indicated, that the AA, AT, GF and NS peptides might maintain their structural integrality *in vivo* inside of yeast and bacteria cells, probably also keeping their functionality (Supplementary Material Table 1). However, KV, KS, RN and LK might disintegrate through the action of enzymes in the environment within minutes, but their antimicrobial activity could be interacting with the membranes outside the cell. Inside mammalian cell lines (*in vitro*), all peptides would be stable for hours (Supplementary Material Table 1). Furthermore, the instability index allowed to classified the AA, NS, RN and AT peptides as “stable” in test tube conditions. Although the GF, KV and LK were classified as “unstable”, their instability index (all values under 53) were not too high above the threshold (40) (Supplementary Material Table 1). In regards of the intestine-like environment stability predictions, none of the peptides were classified with “low” stability. Instead, AA and KS were classified with “high” stability in this proteolytic environment (Supplementary Material Table 1).

The predictions performed with the Peptide Cutter Tool established, that lowly specific enzymes involved in the digestive system functions of vertebrates (Trypsin) or isolated from a fungus (LysN and Proteinase K) or bacteria (LysC and Thermolysin) can degrade all the peptides evaluated in this study (Supplementary Material Table 1). Also, chymotrypsin, a protein isolated from the porcine pancreas gland, and pepsin, a protein isolated from gastric juice of dogs, were predicted to degrade all the peptides except for AA. Furthermore, the inorganic molecule CNBr might possibly fragment KS, NS, AT and GF (Supplementary Material Table 1). In addition, some of these eight new peptides are susceptible to be degraded by proteins isolated from bacteria of clinical importance. For example, RN and GF peptides might be susceptible to Clostripain originating from *Clostridium sp.* and the NS peptide may be fragmented by the Asp-N endopeptidase originating from *Pseudomonas sp.* as well as the Asp-N and Glutamyl endopeptidase and Staphylococcal peptidase I originating from *S. aureus* (Supplementary Material Table 1).

### Proteins and antimicrobial peptides already described have sequence similarities to the eight new peptides: KS, NS, AT, KV and LK might have a possible precursor sequence from proteins involved in gene regulation and all the peptides of this study presented similarities with peptides already described displaying antibacterial activity

Protein alignments with the peptides’ sequences performed with the PIR showed, that the new peptides might have a possible precursor sequence from proteins already described in several organisms and taxonomic groups (Table 2). Three of these proteins have been found to be involved directly or indirectly in regulating gene expression. In addition, the KV and LK peptides’ sequences, as other described peptides like for example Buforin (Park *et al*., 1996), matched with those of histones. Analysis of the taxonomic groups from the proteins matching with the sequence of each peptide showed, that all peptides had a possible precursor sequence from proteins of amphibian origin. On the other hand, the Peptide Match Tool of the APD indicated, that the new peptides of this study had similarities of between 36 to 50 % to already described peptides (Table 2). Surprisingly, the KS peptide sequence is similar to a reported synthetic peptide sequence. All the peptides already described which matched with the eight new peptides of this study showed antibacterial, antifungal, antitumorigenic, cytotoxic, antiparasitic and/or antiviral activity (Table 2).

**Table 2.** Proteins and peptides with similar sequences to the eight new peptides. The PIR and the APD were used to compare the sequences of the peptides used in this study with proteins and peptides already described. All the peptides, except for GF, were found to have a total match with the sequence of proteins according to PIR. The majority of the protein and peptide matches were identified in the amphibian taxonomic group.

| ID | Protein<sup>1</sup> | Peptide<sup>2</sup> | Peptide Source<sup>2</sup> | Peptide Similarity<sup>2</sup> | Peptide Activity<sup>2</sup> |
| --- | --- | --- | --- | --- | --- |
| AA | Decaprenyl-diphosphate synthase | Dominulin B<sup>3</sup> | *Polistes dominulus* (Invertebrates) | 44 % | Anti-Gram + & - |
| KS | Myoneurin | BP100 and L5K5W<sup>4</sup> | Library approach | 36 % | Anti-Gram + & - |
| NS | Nuclear receptor corepressor 2 of (SMART) | Latarcin 3b<sup>5</sup> | *Lachesana tarabaei* (Invertebrates) | 43 % | Anti-Gram + & - and Antifungal |
| RN | Malate dehydrogenase | Alyteserin-2c and 2b and Maximin H1 and H4<sup>6</sup> | *Alytes obstetricans* and *Bombina maxima* (Amphibians) | 42 % | Anti-Gram + & -, Antiviral, Antifungal, Antitumorigenic |
| AT | MGC81605 protein | Lasioglossin LL-1<sup>7</sup> | *Lasioglossum laticeps* (Invertebrates) | 39 % | Anti-Gram + & - and Antitumorigenic |
| GF | - | Temporin-SN3 and SN2<sup>8</sup> | *Hylarana spinulosa* (Invertebrates and Amphibians) | 50 % | Anti-Gram+, Antifungal, Cytotoxic |
| KV | Histone H2A | Maculatin 1.4<sup>9</sup> | *Litoria eucnemis* (Amphibians) | 42 % | Anti-Gram + & - and Antitumorigenic |
| LK | Histone H2A | XPF-St7<sup>10</sup> | *Silurana tropicalis* (Amphibians) | 45 % | Anti-Gram + & -, Antifungal, Antiparasitic and Cytotoxic |

ID = Code given to each peptide according to the first and last amino acid of its sequence. Protein = Protein that exactly matches the sequence of the peptide according to the PIR. Peptide = Most similar peptide found to our new peptide after alignment in the APD. Peptide Source = Source (organism) of the most similar peptide found to our new peptide. Peptide Sim. = Percentage of peptide similarity between our new peptide and the best match found. Peptide activity = Antimicrobial activity reported for the most similar peptide found. 1. PIR, Peptide Match Tool (Chen et al., 2013). 2. APD, APD3: Antimicrobial Peptide Calculator and Predictor (Wang & Wang, 2016). 3. APD ID= AP01639 (Turillazzi et al., 2006). 4. APD ID= AP02671 and AP00143 (Badosa et al., 2007). 5. APD ID= AP01013 (Kozlov et al., 2006). 6. APD ID= AP01463, AP01462, AP00832 and AP00527 (Conlon et al., 2009 and Lai et al., 2002). 7. APD ID= AP01465 (Čeřovský et al., 2009). 8. APD ID= AP02274 and AP02273 (Yang et al., 2013). 9. APD ID= AP00770 (Apponyi et al., 2004). 10. APD ID= AP02292 (Roelants et al., 2013).

### Antigenicity of eight new peptides: KV and LK might interact with antibodies and HLA class I and II

The affinity of the peptides to antibodies was considered according to their antigenic propensity and antigenic determinants. Results showed, that the peptides are potentially antigenic and that GF, KV, and LK have antigenic determinants which might activate B cells (Supplementary Material Table 2). Consequently, the HLA class I binding capacity, taken into account as the peptide – HLA class I complex score to trigger an immune response, also predicted that these last three peptides (GF, KV, and LK) have the highest probability to be immunogenic (Supplementary Material Table 2). On the other hand, the probability of the antigenic presentation of the peptides was calculated with the SYFPEITHI Tool. The results showed that RN, AT, KV and LK of the peptides might be presented by >10 HLA class I alleles, but only KV and LK by HLA class II alleles. Furthermore, the predictions showed, that all peptides except KS might bind to the HLA class I. Regarding the HLA class II, the RN, AT, KV and LK peptides might bind to the molecule with high affinity (Supplementary Material Table 2). Taken together, these results indicate that although all peptides had important predictions about their HLA class I and II binding capacity, the KV and LK peptides were ranked in all variables analyzed as the ones that have the greatest probability to interact with the immune system pathways analyzed.

### Three clinical *S. aureus* isolates are resistant to ampicillin, gentamicin and/or methicillin

In accordance with the expected results, the *S. aureus* strain (ATCC® 25923) was classified as susceptible to all of the antibiotics tested (Figure 1 and Supplementary Material Table 3 and 4). Similarly, the 39413 clinical isolate showed also to be susceptible to all of the antibiotics in the diffusion assay (Supplementary Material Table 3). However, the quantitative methodology demonstrated, that the IC50 and EC50 of the ampicillin were higher in this clinical isolate, although, the concentrations were still too low to indicate that it was ampicillin-resistant (Figure 1 and Supplementary Material Table 4). Regarding the three resistant clinical isolates, all showed different resistance profiles, but none of them was resistant to vancomycin (Figure 1 and Supplementary Material Table 3 and 4). Specifically, the analysis showed that the 34026 isolate only displayed resistance to ampicillin, while the 36055 and 39791 isolate showed resistance to ampicillin, gentamicin and methicilin (Supplementary Material Table 3 and 4 and Figure 1). Taken together, these results classified the 39413 C. I. as non-resistant to the antibiotics tested, the 34026 C. I. as ampicillin resistant and the 36055 and 39791 C. I. as methicilin-resistant *Staphylococcus aureus* (MRSA), multiresistant to ampicillin and gentamicin.

**Fig. 1.**
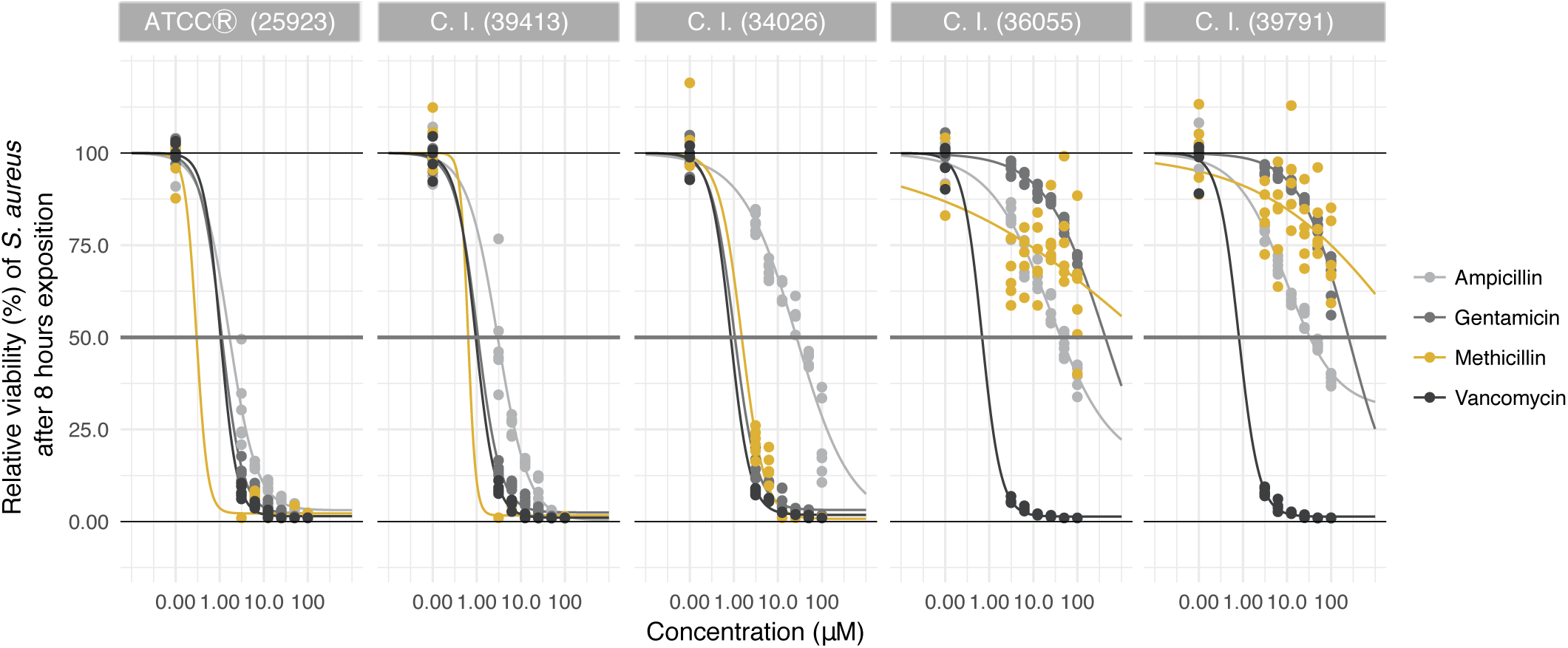
*S. aureus* Clinical isolates (C. I.) resistance profile to four antibiotics. Five *S. aureus*, one of reference (ATCC 25923) and four clinical isolates, were exposed to four antibiotics (ampicillin, gentamicin, methicillin and vancomycin) with the microdilution methodology described above. Their viability was measured after 6, 8 and 24 hours (only viability after 8 hours of exposure results are shown). The black line was calculated as the total growth of the bacteria without exposure to antibiotics. Comparisons with the reference strain were realized with a linear model to determine the C. I. resistances to the antibiotics. The C. I. (39413) was found to be susceptible to all the antibiotics, while the other three (34026, 36055 and 39791) were classified as ampicillin resistant. Furthermore, the 36055 and 39791 C. I. showed to be methicilin resistant *Staphylococcus aureus* (MRSA) and resistant to gentamicin. None of the S. aureus C. I. was identified as vancomycin-resistant *Staphylococcus aureus* (VRSA).

### The peptides individually showed their best activity against *S. aureus* after 24 hours of exposure

Microdilution assays were performed with the peptides individually, the bacterial growth was measured after 6, 8 and 24 hours of exposure and the inhibitory concentration of half the of the bacterial growth (IC50) of the peptides was calculated. Firstly, after 6 hours of exposure, only AT, GF and LK inhibited half of the bacterial growth of a multiresistant MRSA C. I. (Supplementary Material Table 5). After 8 hours of exposure, AT and GF continued inhibiting the multiresistant MRSA C. I., but showed a higher IC50 (Figure 2 and Supplementary Material Table 5). Furthermore, the peptide KV showed to be capable to inhibit the bacterial growth of the multiresistant MRSA C. I. (39791) (Figure 2 and Supplementary Material Table 5).

**Fig. 2.**
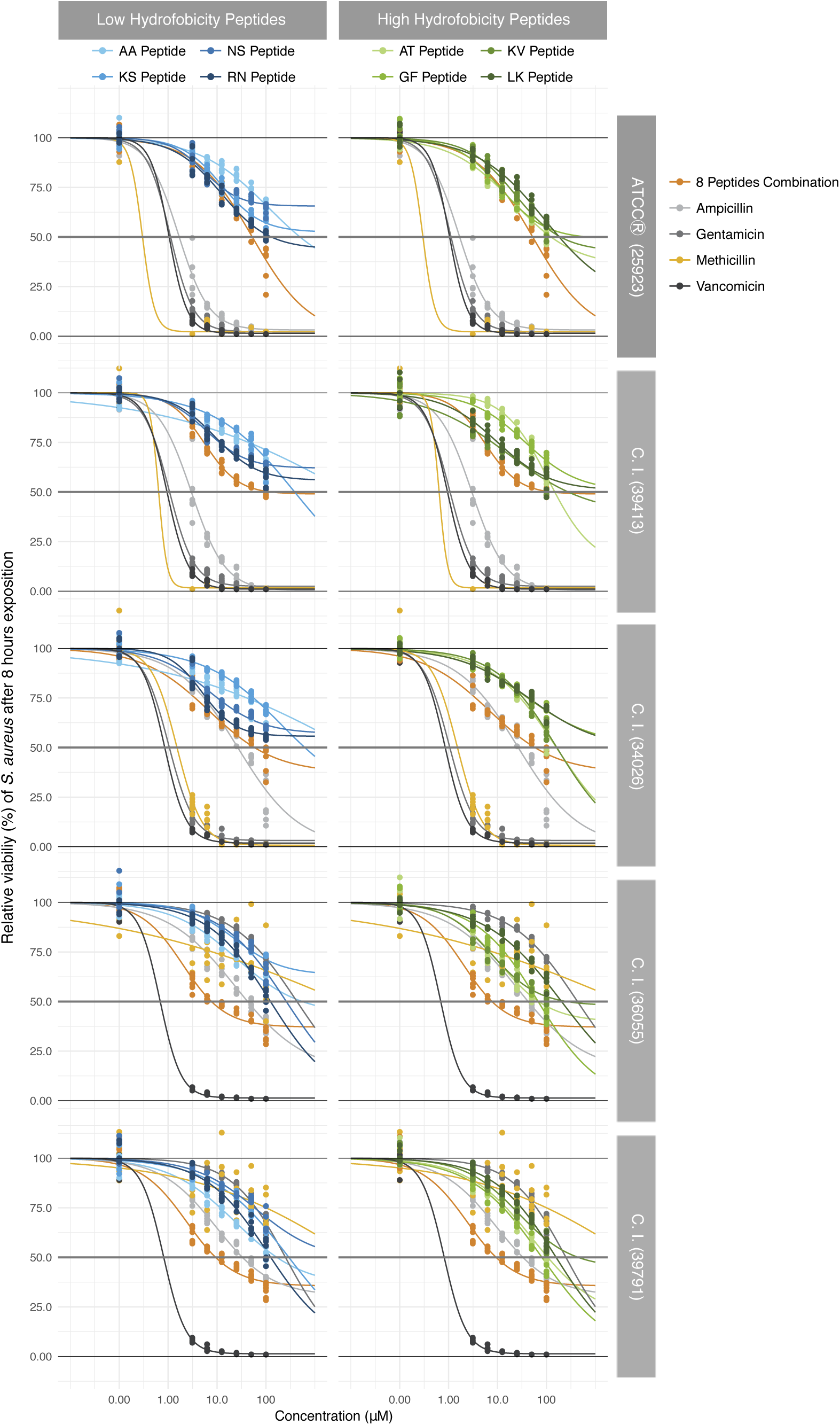
Antibacterial activity of the peptides (individually and combined) against *S. aureus*. Microdilution assays with the peptides individually and combined were performed in order to identify the antibacterial activity of the peptides against a reference *S. aureus* strain (ATCC 25923) and four C. I. *S. aureus* with different antibiotic resistance profiles (39413, 34026, 36055 and 39791). The bacteria viability was measured after 6, 8 and 24 hours of exposure (Only the 8-hour exposure viability results are shown). The individual results for each peptide were separated according to their hydrophobicity. High hydrophobicity individual peptides were more potent inhibiting bacterial growth than low hydrophobicity individual peptides. The eight peptide combination was more potent than the peptides individually and it was able to inhibit bacterial growth of four of the *S. aureus* evaluated. Furthermore, this antibacterial activity was stronger in the multiresistant clinical isolates (36055 and 39791).

Most of the peptides strongest antibacterial activity against the *S. aureus* bacteria was reached after 24 hours of exposure (Supplementary Material Table 5). For example, AA, KS and RN inhibited the *S. aureus* bacteria after 24 hours of exposure, a capacity that was not displayed in the shorter exposure. Furthermore, GF inhibited the reference strain growth at the lowest IC50 found between the results of the individual peptide (22.8μM) (Supplementary Material Table 5). Although overall, none of these peptides individually inhibited the bacteria growth of all the bacteria tested, AA, AT and KS peptides had the capacity to inhibit the bacteria growth of at least two of *S. aureus* (Supplementary Material Table 5). On the other hand, in general, the most susceptible bacteria to the peptides were the reference strain (ATCC 25923) and, surprisingly, the clinical isolate 36055 (Supplementary Material Table 5). Also, the only peptide which did not show antibacterial activity against the *S. Aureus* tested at any time was the NS peptide.

In addition, statistical analysis of the antibacterial activity of the peptides showed that after 24 hours of exposure, the capacity of the peptides to inhibit the bacterial growth was significantly reduced, depending on the C. I. it was exposed to when comparing with the reference strain (Supplementary Material Table 6). For example, the GF peptide’s antibacterial activity decreased when it was exposed to all the C. I. tested. However, the NS and RN peptides’ antibacterial activity did not decrease against any of the C. I. exposed to (Supplementary Material Table 6). Interestingly, taken together, these results showed that the C. I. 39413 growth was significantly less inhibited by the peptides when comparing with the reference strain (Supplementary Material Table 5). Still, these decreases in the antibacterial activity were not comparable to those of the antibiotics when exposed to resistant bacteria. Finally, the antibacterial activity of the individual peptides were compared with the antibiotics. Results showed that overall, the antibiotics decreased the growth of all the *S. aureus* significantly more than the peptides individually (Supplementary Material Table 7).

### The peptides combined showed their best activity against the MRSA *S. aureus* after 8 hours of exposure

In consequence of the low antibacterial activity that the individual peptides exhibited, antibacterial assays were performed with all the peptides combined. Results showed that the combination had a significantly more potent activity against the bacteria. Consequently, the combination exhibited the capacity to inhibit four of the *S. aureus* and in three of them, this capacity was shown after 24 hours of exposure. Specifically, after 6 hours of exposure, the eight peptide combination inhibited the bacterial growth of the reference strain and the two C. I. multiresistant MRSA (Supplementary Material Table 8). Then, after 8 hours of exposure, the combination showed its broader antibacterial activity inhibiting the bacterial growth of the reference strain and three C. I. with a very low IC50 (Figure 2 and Supplementary Material Table 8). Finally, after 24 hours of exposure, the eight peptide combination had the capacity to keep inhibiting the bacterial growth of the reference strain (ATCC® 25923) and the two C. I. multiresistant MRSA (Supplementary Material Table 8). In addition, this antibacterial activity did not decrease in a significant way when the combination was exposed to C.I. when comparing with the reference strain. Nevertheless, the combination was not able to inhibit the bacterial growth of the C. I. non-resistant 39413.

The peptides (combined) and antibiotics antibacterial activity was compared. Interestingly, all the antibiotics inhibit both the bacterial growth of the reference strain and the non-resistant C. I. significantly more than the eight peptide combination. Similarly, gentamicin, methicilin and vancomycin inhibited the bacterial growth of the C. I. ampicillin-resistant significantly more than the peptide combination and vancomycin had the capacity to inhibit the bacterial growth of the multiresistant MRSA C. I. more significantly than the peptide combination (Supplementary Material Table 8). However, the eight peptide combination was able to inhibit the bacterial growth of these more effectively than ampicillin, gentamicin, and methicilin (Supplementary Material Table 8).

### The eight peptide combination and the AT and LK peptides individually displayed cytotoxic activity only against THP-1 monocytes

The overall results indicate significant differences between the cell culture types (Figure 3 and Supplementary Material Table 9). Firstly, assays performed in the VERO cell line showed, that the peptides did not have cytotoxic activity against these fibroblasts, neither individually nor combined. In contrast, the antibiotics evaluated (ampicillin, gentamicin and vancomycin) negatively affected the viability of the cells, even though none decreased the cells viability more than 50 % (Figure 4 and Supplementary Material Table 9). The positive control, doxorubicin, inhibited the cells growth significantly and presented an IC50=56.71μM as expected (Figure 4 and Supplementary Material Table 9). Contrary to the above results, cytotoxic activity assays performed in the THP-1 monocytes cell culture showed, that all high hydrophobicity peptides and the eight peptide combination inhibited the cells viability in a significant way. However, only AT and LK individually and the eight peptide combination inhibited more than the 50 % of the viability of the cells between the concentrations tested (Figure 4 and Supplementary Material Table 9). In addition, the antibiotics evaluated did not inhibit the monocytes viability (Figure 4 and Supplementary Material Table 9). However, the positive control, doxorubicin, significantly inhibited the THP-1 cells growth and presented a lower IC50 (Figure 4 and Supplementary Material Table 9) than the one calculated for the VERO cell line.

**Fig. 3.**
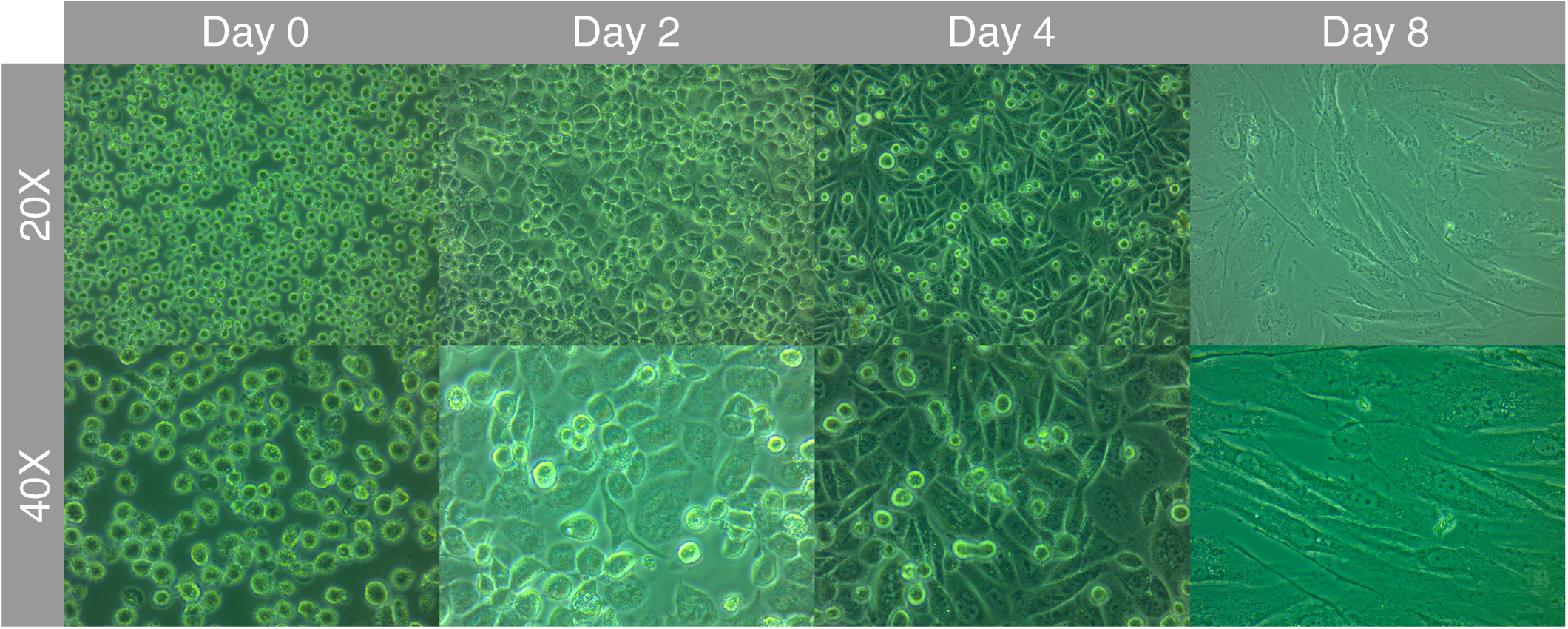
THP-1 monocytes differentiation into macrophages after 200 nM PMA exposure. THP-1 monocytes were exposed to 200 nM PMA for three days, followed by five days of incubation in RPMI media with supplements free of PMA. Photos taken at day 0 of the THP-1 monocytes before PMA treatment show their suspension phenotype and small size. At day two cells started to change their morphology and to adhere. After four days, macrophages elongated and remained in their adherent phenotype. After eight days, macrophages were fully differentiated.

**Fig. 4.**
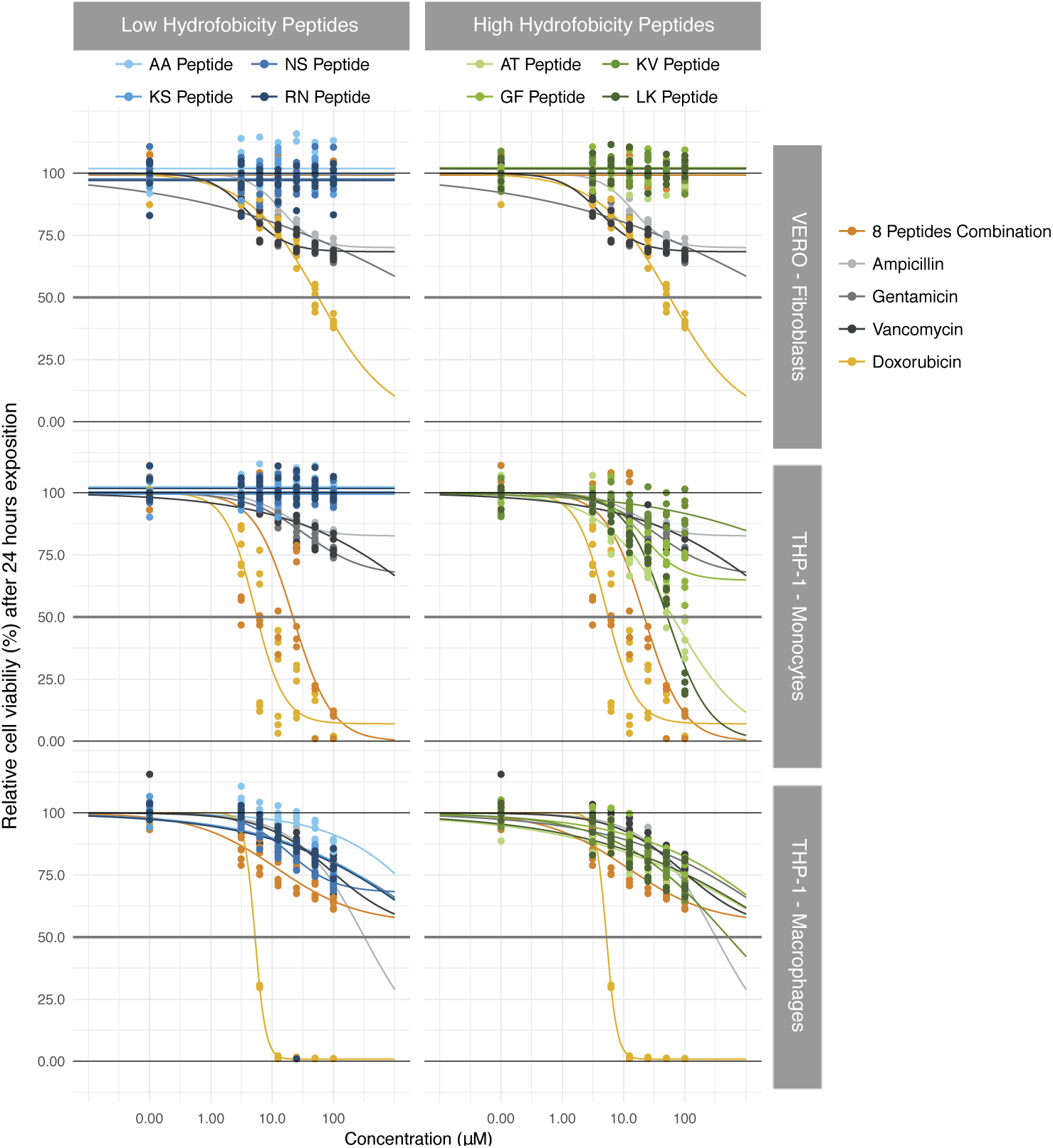
Cytotoxic activity of eight new peptides (individually and combined) in three cell lines. The peptides’ effect (individually and combined) on the viability of the cells was evaluated in three cell types (VERO-fibroblasts, THP-1-monocytes and THP-1-macrophages) via MTT assays. Cytotoxicity assays presented significant differences between cell lines. The individual peptides and the eight peptide combination did not show cytotoxic activity in the VERO – fibroblasts nor in THP-1 macrophages. In contrast, AT, LK peptides and the eight peptide combination showed cytotoxic activity against THP-1 monocytes.

Ultimately, THP-1 monocytes were differentiated into macrophages using PMA. Cells rapidly changed their suspension phenotype typical for monocytes to the adherent expected phenotype of macrophages, further morphological and size changes were observed (Figure 3). In addition, in order to confirm the differentiation, the expression of recognized macrophages markers (CD11b, CD14 and CD36) was analyzed. As expected, the results showed that macrophages expressed the CD11b marker more than monocytes, while the CD14 and CD36 markers were expressed in both macrophages and monocytes. Following, we assessed cytotoxic activity assays with the THP-1 macrophages as described previously. Results showed that all the peptides inhibited less than 50 % of the viability of the macrophages (Figure 4 and Supplementary Material Table 9). In contrast with the previous results in the monocyte cell line, antibiotics inhibited the viability of the cells, but still, none were able to decrease the 50 % of the viability of the cells (Figure 4 and Supplementary Material Table 9). Finally, consistent with the monocyte assays results, doxorubicin decreased the viability of the macrophages significantly and presented a similar IC50 (Figure 4 and Supplementary Material Table 9) to the one calculated for the THP-1 monocytes.

### Each evaluated peptide (AA, GF and RN) displayed dose-dependent immunomodulatory activity

After differentiating the THP-1 monocytes to macrophages as mentioned previously, the macrophages were exposed to three concentrations (0, 6.25, 25 μM) of three peptides selected for their high antibacterial and low cytotoxic activity (AA, GF and RN). Furthermore, experiments were assessed in three different stimulation scenarios: without stimulation, with *E. coli* LPS or with *S. aureus* lysate. In the supernatants of the exposed cells, the concentration of 13 cytokines (IL-1β, IFN-α, IFN-γ, TNF-*α*, CCL2 (MCP1), IL-6, IL-8, IL-10, IL-12p70, IL-17A, IL-18, IL-23 e IL-33) was determined with the LEGENDplex™ Human Inflammation Panel kit 13-plex (Biolegend). The results obtained indicated that the peptides could modulate the secretion of four (IL-1β, CCL2, TNF-*α* and IL-8) of the 13 cytokines analyzed.

A Bartlett’s test was performed to identify if the cytokines secretion variables were correlated. Results indicated a strong correlation between the variation of the variables, hence a discriminal function analysis (DFA) was performed to determinate whether each peptide modulated the cytokine secretions in a significantly different manner (Figure 5). A DFA was performed for each different stimulation scenario, results demonstrated that each peptide modulated the cytokine secretion differently. Specifically, in the non-stimulation scenario (Figure 5), the discriminant function Can 1 explained the 95 % of the variation between the variables related to cytokine secretion, were the AA peptide increased the secretion of IL-8 and decreased that of the IL-1β, CCL2 and TNF-*α* cytokines. On the contrary, while both the GF and RN peptides decreased the secretion of IL-8, only GF increased the secretion of TNF-*α* and IL-1β and only RN increased the secretion of CCL2. Regarding the *E. coli* LPS stimulation scenario (Figure 5) the discriminant function Can 1 explained 72.7 % of the variation, while the Can 2 explained 27.3 %. In this analysis, the RN peptide increased all the cytokines secretions, while the AA and GF peptides decreased the cytokine secretions, especially IL-1β/TNF-*α* and CCL2/IL-8 respectively. Finally, the *S. aureus* lysate stimulation scenario (Figure 5) showed, that the peptides modulated the cytokine secretions in an again different manner. The discriminant function Can 1 explained 97 % of the variation, where the AA peptide increased the IL-8 and CCL2 cytokine secretion, while RN and GF decreased these cytokine secretions. In addition, in this scenario, the RN peptide also decreased the IL-1β and TNF-*α* secretion, while the GF peptide increased their secretion.

**Fig. 5.**
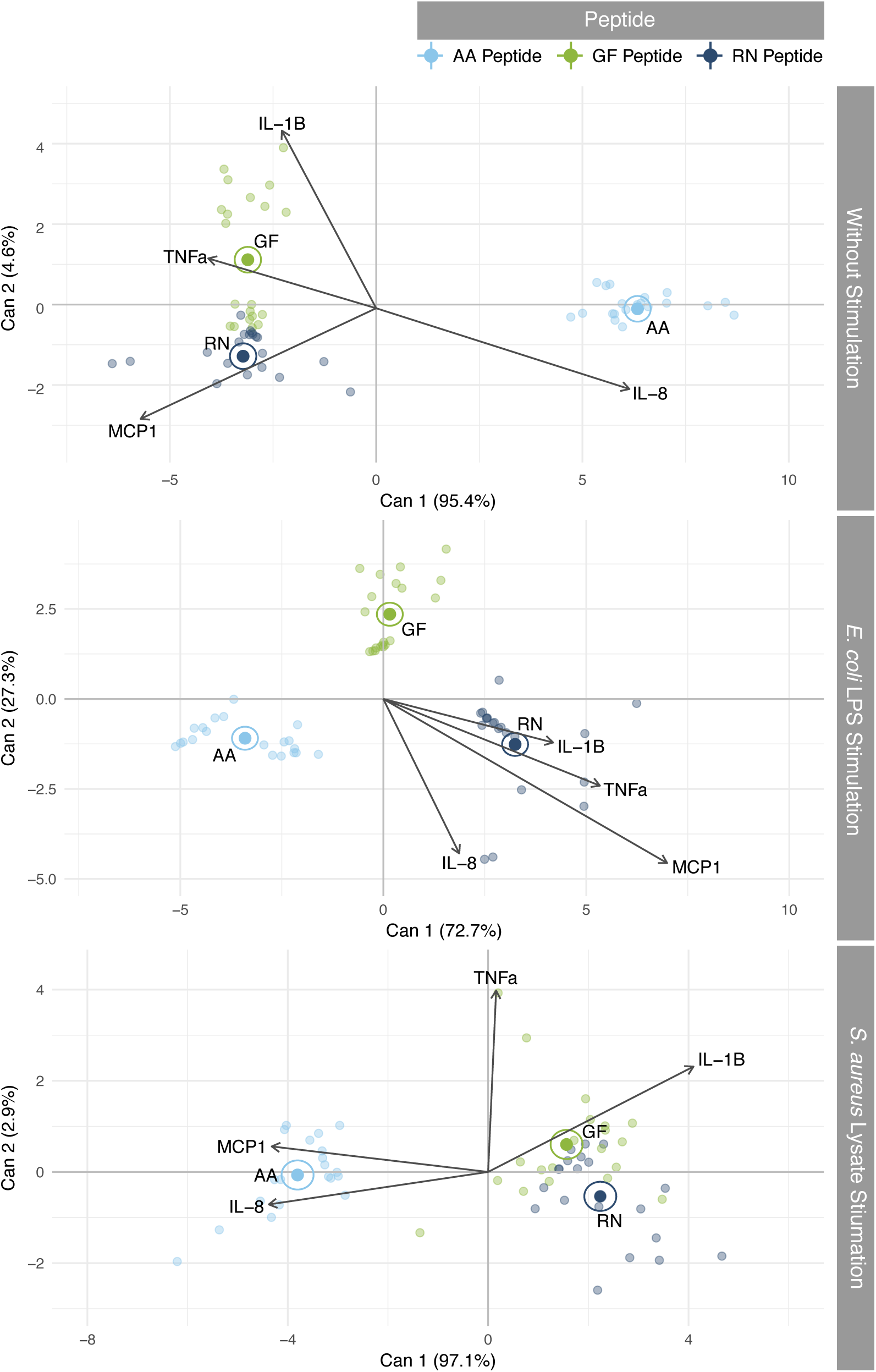
DFA analysis of the immunomodulatory activity of three peptides in three stimulation scenarios. The immunomodulatory activity of the peptides individually was evaluated with the LEGENDplex™ Human Inflammation Panel (13-plex) kit (Biolegend). THP-1 macrophages were exposed to the peptides in three stimulation scenarios (non-stimulation, stimulation with *E. coli* LPS and stimulation with *S. aureus* lysate) and the concentration of the cytokines secreted to the culture media was calculated with flow cytometry. Results of four cytokines (IL-1β, CCL2 (MCP1), TNF-*α* and IL-8) were analyzed via DFA. Depending on each stimulation scenario, the peptides modulate the cytokine secretion to the media. Furthermore, this cytokine secretion modulation showed significant differences between the peptides.

To determine the existence of significant differences in the effect the peptides had on each of the cytokines secretions between the two peptide concentrations tested, a model selection with the Akaike information criterion was performed. In all the cases, the best linear model was the one that considered the peptide, the peptide concentration, and the stimulation scenario. Then, a linear model analysis was performed with each cytokine as the response variable. The results showed, that IL-1β secretion, without the presence of the peptides, significantly increased when the macrophages were stimulated with the *E. coli* LPS, while there were no differences in IL-1β secretion when the macrophages were stimulated with the *S. aureus* lysate. When the macrophages were exposed to the AA and GF peptide in the non-stimulation scenario, the cytokine secretion increased at the lowest peptide concentration. In addition, we observed an increment in this effect, especially when the GF peptide concentration was higher. In contrast, at the low concentration, RN peptide increased the IL-1β secretion significantly more, while there was no observable modulation of the cytokine secretion at the high concentration (Figure 6 and Supplementary Material Table 10). In the *E. coli* LPS stimulation scenario, the three peptides decreased the IL-1β secretion at their lowest concentration, but then at the highest concentration, they significantly increased the cytokine secretion, especially the RN peptide (Figure 6 and Supplementary Material Table 10). Finally, when the macrophages were stimulated by the *S. aureus* lysate, the three peptides increased the IL-1β secretion at low concentrations. Furthermore, when the GF and RN concentration increased, the concentration of the IL-1β further increased, while it remained similar when the AA concentration increased (Figure 6 and Supplementary Material Table 10).

**Fig. 6.**
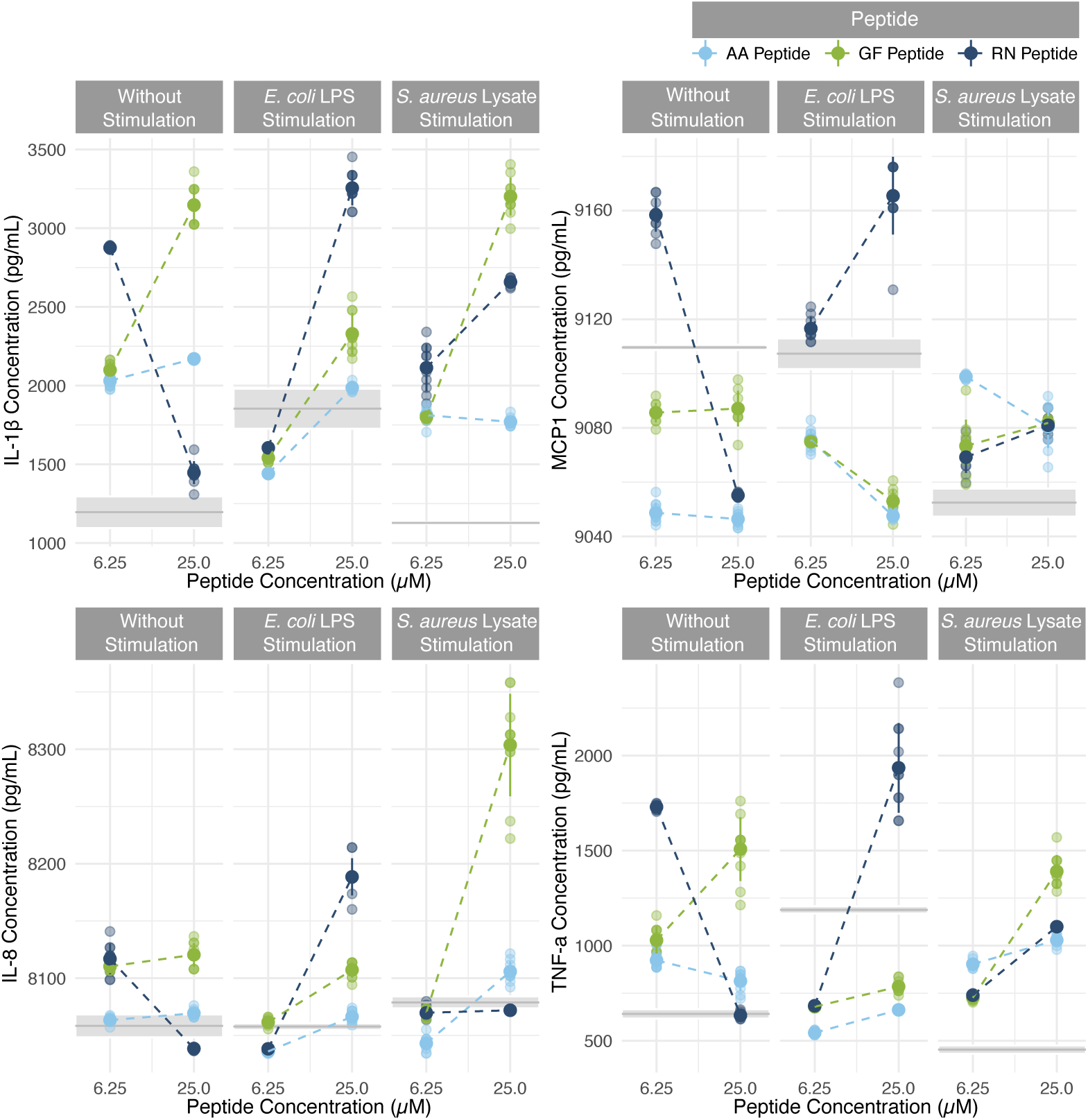
Analysis of the effect of peptide concentration on the secretion of four cytokines. To determine if the concentration of the peptides had an effect on the secretion of each of the four cytokines (IL-1β, CCL2 (MCP1), IL-8 and TNF-*α*), a model selection was performed using the Akaike information criterion, followed by a linear model analysis. The concentration of the peptides significantly affects the cytokine secretions, and this effect further depends on the stimulation scenario. The grey line and it’s corresponding standard deviation represents the reference cytokine concentration for each stimulation scenario. Each treatment consists of two peptide concentrations (6.25µM and 25µM), the lightly coloured dots represent the raw data, the bigger dots represent the mean and the error bars it’s corresponding standard deviation.

The CCL2 chemokine concentrations in the non-stimulation and the *E. coli* LPS stimulation scenarios were not significantly different, while they decreased significantly in the *S. aureus* lysate stimulation scenario. When the macrophages were exposed to the AA and GF peptides in the non-stimulation scenario, the CCL2 secretion decreased, and this effect did not change when the concentration of the peptides increased. In contrast, when the cells were exposed to the low concentration of the RN peptide, the chemokine secretion increased, but when they were exposed to its high concentration, the chemokine secretion decreased (Figure 6 and Supplementary Material Table 10). In the *E. coli* LPS scenario, the AA and GF peptides kept decreasing the CCL2 secretion, and in this case, this effect augmented when the peptides concentration increased. On the contrary, exposing macrophages to the RN peptide at a low concentration did not have any effect on the chemokine secretion, while the peptide at a high concentration increased its secretion (Figure 6 and Supplementary Material Table 10). In the *S. aureus* lysate stimulation scenario, all the peptides increased the CCL2 secretion in their low concentration, but this effect decreased when the AA peptide was presented at a higher concentration, while it remained stable when the GF and RN concentration increased (Figure 6 and Supplementary Material Table 10).

Regarding the IL-8 cytokine concentration, our results show that after the macrophages were not stimulated and when they were stimulated with *E. coli* LPS and *S. aureus* lysate, there were no significant differences in the cytokine secretion and with *S. aureus* lysate. However, when the macrophages were not stimulated but exposed to the peptides, the low concentration of the GF and RN peptides increased the IL-8 secretion and remained unaffected when the RN concentration increased, but it decreased when the GF concentration increased (Figure 6 and Supplementary Material Table 10). In the *E. coli* stimulation scenario, the three peptides slightly decreased the IL-8 at their low concentration and increased the cytokine secretion in their high concentration. Similarly, in the *S. aureus* stimulation scenario, the three peptides marginally decreased the cytokine concentration and increased it at their high concentration, especially the GF peptide (Figure 6 and Supplementary Material Table 10).

Finally, when the macrophages were stimulated with the *E. coli* LPS, the TNF-*α* concentration significantly increased compared to the non-stimulation scenario, while in the *S. aureus* lysate stimulation scenario the cytokine concentration decreased. When the macrophages were not stimulated, the exposure to a low concentration of the peptides increased the TNF-*α* concentration, but exposed to a high peptide concentration, both the RN and the AA peptides decreased the cytokine secretion, while it was further increased by the GF peptide (Figure 6 and Supplementary Material Table 10). On the other hand, in the *E. coli* LPS stimulation scenario at a low concentration exposure, all the peptides decreased the TNF-*α* concentration and increased it at their high concentration, especially the RN peptide. Finally, when the macrophages were stimulated with the *S. aureus* lysate, the peptides at a low concentration increased the TNF-*α* concentration, and this effect further increased when the peptides were presented at a higher concentration (Figure 6 and Supplementary Material Table 10).

## Discussion

The need to search for new alternatives against multiresistant infectious diseases comprehend not only new antibiotic agents but also new therapeutic agents for alternative therapies such as immunomodulatory therapy. With the objective of controlling *S. aureus* infections, it has been proposed that treatment should include immunomodulatory therapy (Wardenburg, Williams Missiakas, 2006 and Thangamani, Younis Seleem, 2015). Therefore, as mentioned previously, this study aimed to identify the potential of eight new peptides as antibacterial and immunomodulatory agents as a basis for the search of new therapeutic agents for the topic prevention and treatment against *S. aureus. In silico* analysis of the antimicrobial activity, stability, sequence similarities and antigenicity of the eight new peptides features were performed. Then, the antibacterial activity of the eight peptides individually and combined was evaluated. Cytotoxicity assays in three cell lines, including two of immune-related cells, were performed. Those results, together with the antibacterial activity of the peptides, allowed to select AA, GF and RN for the immunomodulatory activity assays. Finally, the evaluation of the capacity of the peptides to stimulate and/or inhibit the production of cytokines were performed in three different simulation scenarios.

*In silico* results determined that, although the eight peptides described in this study do not share the same characteristics, all predictions regarding their general characteristics indicated their potential to be antimicrobial peptides. The predictions about their stability features suggested the possible environments where they would be active as well as the ones where they could easily degrade. Positive net charge, short length, amphipathic composition and formation of an *α* helix are common characteristics presented by host defence peptides with antimicrobial activity, especially because these features are associated with the capacity of the peptides to interact with living cell membranes (Zhang Gallo, 2016). This interaction is important because most of the mechanisms of action of the antimicrobial peptides described are focused on the capacity of the peptide to disrupt or translocate living membranes (Nguyen, Haney Vogel, 2011). Hence, these characteristics are usually correlated with the biological activity of the peptides (Soltani, Keymanesh, Sardari, 2007). We also evaluated *in silico* the stability of the peptides in certain environments. A major limitation of the peptides biological activity is their susceptibility to proteases (Meng Kumar, 2007). However, according to the *in silico* analysis of the peptides in this study, three would be considered highly stable in milieu proteolytic environments. Nevertheless, all the peptides were determined to be susceptible to a wide range of proteases, indicating an important limitation of the peptides. Studies have proposed alternative modifications to increase the stability of antimicrobial peptides in proteolytic environments, such as the amidation and acetylation of the terminals or by introducing tryptophan substitutions at cleavage sites (Strömstedt *et al*., 2009). These alternatives should be taken into consideration for further studies with these peptides.

Alignments *in silico* of the peptides’ sequences found proteins and antimicrobial peptides already described with sequence similarities to the eight new peptides of this study. Previous studies have found, that many antimicrobial peptides are derived of the cleavage of proteins (Stewart *et al*., 2008). For example, Buforin I, a potent antimicrobial peptide, is the product of the cleavage of the Histone H2A by pepsins (Kim *et al*., 2000). Similarly, LK and KV peptides were found to have sequence similarities with histones. These results might predict seven possible protein precursors of the eight new frog peptides described in this study, an assumption supported by the fact that the majority of these possible protein precursors were identified in the taxonomic group of amphibians. On the other hand, all the eight new peptides were identified to have sequence similarities with antimicrobial peptides already described. Some studies have considered, that many peptides are derived from other larger peptides. In some cases, the shorter new peptides present a more potent biological activity (Zhao *et al*., 2016). In addition, these sequence similarities allow classifying the peptides into the different families of antimicrobial peptides already described (Khamis *et al*., 2014), however further studies would be needed to ascertain the pertinence of each peptide to the families that are suggested here.

Immunogenicity is one of the biggest concerns in the search for new antimicrobial agents (Yeaman, Yount, 2003). As peptides are amino acid sequences of short length, they can be presented via HLA class I and class II and by antibodies. Hence, in this study, we performed analysis with bioinformatic tools, which are commonly used to identify antigens that might be recognized by the immune system, along with specific tools that directly predict the antigenicity of the peptides. The *in silico* analysis determined, that LK and KV peptides were predicted to have a major probability to trigger an immune response. Nonetheless, AA, KS and GF peptides were considered too short to be presented by the HLA class I and II, which is considered a major advantage for their subsequent use as therapeutic agents (Fox, 2013).

*S. aureus* infections are one of the major concerns world-wide due to their increased resistance to antibiotics, especially because they can cause several severe health issues when they are acquired in healthcare environments (Fournier Philpott, 2005). Here, we evaluated the antibacterial activity of the peptides against a reference strain and four clinical isolates, including two methicillin-resistant *Staphylococcus aureus* (MRSA). The results with the MRSA bacteria indicated, that the use of the peptides instead of antibiotics significantly decreased the bacterial growth more, highlighting the importance of evaluating the new antibacterial agent candidates against multiresistant bacteria. Although the peptides individually decreased the bacterial growth, the IC50 of their antibacterial activity ranges between 20-100μM. These results contrast with other synthetic peptides, which with a calculated IC50 range of in-between 4μM and 8μM, have shown a much more potent antibacterial activity against *S. aureus* (Mohamed, Abdelkhalek Seleem, 2016).

Interestingly, among the antibacterial assays performed with the peptides individually, the GF peptide showed the best results against one of the MRSA, AT the most prolonged activity and AA the best antibacterial activity against several *S. aureus*. Meanwhile, the *in silico* results determined, that the eight new peptides have sequence similarities with described peptides reported to have antibacterial activity. Impressively, the peptides that matched the sequence of AT, GF and KV were described in other studies as highly active against reference strains of *S. aureus* (Čeřovský *et al*., 2009, Yang, *et al*., 2013 and Apponyi *et al*., 2004). Although highly homologous peptides can greatly vary in their biological activity, peptide families usually show an overall antimicrobial activity addressed against a specific group of bacteria. This is the case of the Temporin SN3 and SN2 peptides, which showed a 50 % of sequence similarity with the GF peptide, and belong to the Gram + antibacterial peptide family of Temporins (Conlon, 2011). On the other hand, the NS peptide was the only peptide that did not present antibacterial activity against any evaluated *S. aureus*. We consider that, as shown in the *in silico* results, the susceptibility of the peptide to the *S. aureus* staphylococcal peptidases degraded it during the bacterial exposure in the assays.

In consequence of the variation in the antibacterial activity shown by the individual peptides, antibacterial assays were assessed with all the eight peptides combined. It has been proposed, that this increased activity might be a consequence of the synergic action of the peptides, a suggested common phenomenon in antimicrobial peptide interactions (Yu, 2016). Nonetheless, the combination of molecules with biological activity can result in synergistic, additive or antagonistic interactions (Yu, 2016). Therefore, further studies should be conducted with the eight peptides combined, in order to understand their action mechanisms and to sort out irrelevant combinations of peptides in order to find combinations enhancing the desired antibacterial activity.

In this study, the peptides decreased the viability of the THP-1 macrophages more than the viability of the THP-1 monocytes. Accordingly, it has been reported that the monocytes are especially susceptible to apoptosis and are easily stimulated to apoptosis, while macrophages are considered to be resistant to many apoptotic stimuli (Liu *et al*., 2001). On the other hand, the *in silico* analysis identified that AT has a sequence similar to Lasioglossin LL-I, a peptide reported to have antitumorigenic activity in a wide range of cancer cell lines, including the leukaemia THP-1 cell line used in this study (Čeřovský *et al*., 2009). Meanwhile, antitumorigenic activity assays have not been performed with the peptide that has a similar sequence to the LK peptide, XPF-St7, but it was reported to have cytotoxic activity (Roelants *et al*., 2013). Hence, we propose that as two peptides (AT and LK) and the eight peptide combination showed potent cytotoxic activity against the THP-1 cell line, further studies could be of interest in a subsequence analysis as antileukemic agents. Nevertheless, because of their cytotoxic activity and despite the antibacterial activity of these peptides and the peptide combination found, AT, LK and the eight peptide combination were discarded for the immunomodulatory activity assays.

Regarding the immunomodulatory activity, this study evaluated the capacity of three peptides to modulate the macrophages secretion of a wide cytokine panel. Similar studies evaluating a smaller number of cytokines have found, that some peptides can modulate the macrophages’ cytokines secretion and that this activity is usually described as a complex effect in different cytokines (Pantic *et al*., 2014). For example, Tigerinin 1R, Tigerinin 1V and Tigerinin 1M demonstrated to have the capacity to increase the production of the IL-10 anti-inflammatory cytokines and to decrease the IFN-y secretion in a mouse macrophage cell line (Pantic *et al*., 2014). Other studies have shown, that peptides can increase the production of pro-inflammatory cytokines, such as Frenatin 2D (TNF-*α*, IL-1β and IL-12) (Conlon *et al*., 2013) and Escuelin-2CHA (TNF-*α*) (Attoub *et al*., 2013). Concomitant to this results, each peptide tested (AA, GF and RN) showed a different profile of increasing or decreasing the cytokine secretions by THP-1 macrophages. Interestingly, these profiles varied according to the stimulation scenario which the macrophages were exposed to. These results might suggest the existence of some specificity of the peptides to modulate the immune response depending on the microorganism to which they are exposed to, or to which the macrophages respond. Studies have proposed that this immunomodulatory activity of the antimicrobial peptides found in frogs might indicate, that the peptides also regulate the immune response in terms of pro- and anti-inflammatory cytokine secretions (Pantic *et al*., 2014, Conlon *et al*., 2011 y Conlon *et al*., 2014).

In regards to this study, interestingly all the peptides evaluated, but especially the GF peptide, increased the cytokine secretion (IL-1β, CCL2, TNF-*α* and IL-8) when the macrophages were stimulated with *S. aureus* lysate. Studies have determined, that the IL-1β secretion by bone-marrow-derived cells, such as macrophages, is required for neutrophil recruitment and bacterial clearance in *S. aureus* infections (Miller *et al*., 2007). The increase in the IL-8 concentration allows neutrophils to significantly reduce the bacterial load. In addition, an increase in this cytokine decrease the neutrophils apoptosis that is usually enhanced by *S. aureus* (Zurek, Pallister Voyich, 2015). However, as *S. aureus* induces inflammatory cytokines and causes inflammatory diseases, during infection the CCL2 and TNF-*α* concentrations are commonly increased during infection, a further increment in these cytokines’ secretion would exacerbate the infection. The aforementioned and several other studies have demonstrated, that cytokines play an important role in *S. aureus* clearance and infections healing. This has been confirmed in recombinant cytokine treatments *in vivo* in mammals studies, were the enhancement of IL-1β concentrations successfully cured the infections in 75 % of the cases (Daley *et al*., 1993). Therefore, new agents capable to modulate the cytokine secretion in a *S. aureus* infection scenario are of medical interest for developing new therapies.

## Conclusions

The worldwide emergence of multi-antibiotic resistant infectious diseases, as the ones caused by *S. aureus*, frame the need to search for new alternatives for therapeutic agents. As mentioned previously, this study aimed to identify the potential of eight new peptides as immunomodulatory or antibacterial agents, or both, as a basis for the search of new agents for the topic prevention and treatment against *S. aureus*. With this intention, *in silico* analysis, antibacterial and cytotoxic assays and the evaluation of the capacity of the peptides to stimulate and/or inhibit the production of cytokines were assessed. The analysis identified AA, GF and AT as the peptides with best antibacterial performance individually against the *S. aureus* evaluated. However, the combination of the eight peptides was the better treatment to decrease the bacterial growth, especially of the clinical isolates. Furthermore, the majority of the peptides, except for LK, AT and the peptide combination, did not showed cytotoxic activity, which is a desirable feature for their subsequent use as drugs. Nevertheless, LK and AT peptides are proposed as molecules of interest for the study of new anti-leukaemia agents. Finally, AA, GF and RN peptides presented immunomodulatory activity in THP-1 macrophages. This feature depended on the peptide and the stimulation which the macrophages were exposed to. Results indicate that AA, GF and RN might be of interest for immunomodulatory therapy against *S. aureus* infections. Nevertheless, further studies should be performed to better understand the peptide interactions with the immune system.

## Methodology

### Peptide Synthesis

Eight new antimicrobial peptides previously described in the Human Genetics Laboratory of the Los Andes University in Bogotá, Colombia, named after their first and last amino acid of their sequence (AA, KS, NS, RN, AT, GF, KV and LK) were used in this study. Their synthesis was performed by GL Biochem Ltd. (Shangai, China) via the solid phase method. Afterwards, the analysis and purification of the peptides were performed by reversed-phase HPLC using a Venusil XBP-C18 RP-HPLC column (4.6mm × 250mm) at a flow rate of 1mL/min and a linear gradient of 20:80 % of acetonitrile:water with 0.1 % trifluoroacetic acid with detection of 250 nm. Finally, electrospray mass spectrometry was used to corroborate the purity (>98 %) and for the identification of the synthetic peptides.

### *In silico* characterization of eight new peptides individually

General peptide characteristics, stability related features, sequence similarities with previously described proteins and antimicrobial peptides, as well as the antigenicity of the eight new peptides were analyzed with a broad repertoire of databases and bioinformatic tools. Firstly, the “APD3: Antimicrobial Peptide Calculator and Predictor” tool (Wang Wang, 2016) of the Antimicrobial Peptide Database (APD) was used to identify the following general characteristics of antimicrobial peptides in the eight new peptides: Net charge, hydrophobic and hydropathy (GRAVY) ratio, potential capacity to attach to lipid bilayers and other proteins (W-W hydrophobicity and Boman Index) and special features like potential for *α* helices formation or sequence richness in some specific amino-acids. The ProtParam Tool (Gasteiger *et al*., 2005) was used to identify the stability features of the eight new peptides, allowing recognition of their suitability to remain stable in specific environments. Based on the N-terminal amino acid of each peptide, the half-life *In vitro* and *In vivo* was predicted along with the instability and aliphatic index and finally classification of the stability of the peptide. Likewise, the “HLP: Web server for predicting half-life of peptides in intestine-like environment” (Sharma *et al*., 2014) permitted the prediction of the half-life and stability of the new peptides in an intestine-like, proteolytic, environment.

Alignments of the eight new peptides with the Protein Information Resource (PIR) and the APD (Chen *et al*., 2013 Wang, 2016) were developed to predict the possible precursor sequences of the new peptides and to identify the characteristics as well as the antimicrobial activity of similar described peptides. Finally, the antigenicity of the new peptides was analysed *in silico*. The “Predicted Antigenic Peptide Tool” (Reche, 2002) was used to determine the potential binding capacity to antibodies. Furthermore, the possible antigenic presentation of the peptides, by the MHC class I and II pathways, was identified using the SYFPEITHI Tool (Rammensee, 1999). Additionally, the antigenicity (Calis *et al*., 2013) and affinity to class I and II MHC were calculated using the Immune Epitope Database and Analysis Resource (IEDB). The MHC I binding predictions were made on April 17th, 2017 using the IEDB Analysis Resource Consensus Tool for MHC I (Kim *et al*., 2012), which combines predictions from AMM (Nielsen *et al*., 2003 and Lundegaard *et al*., 2008), SMM (Peters Sette, 2005) and comblib (Sidney *et al*., 2008). The MHC II binding predictions were made on the same date and using the IECB Analysis Resource Consensus Tool for MHC II (Wang *et al*., 2008, Wang *et al*., 2010 and Sturniolo *et al*., 1999).

### Determination of the antibacterial activity of eight new peptides individually and combined against a reference strain, a non-resistant clinical isolate and three resistant clinical isolates of *S. aureus*

A reference strain (ATCC 25923), a non-resistant clinical isolate (39413) and three resistant clinical isolates (34026, 36055, 39791) were used in this study. To ascertain the integrity of the bacteria resistance profile, antibiotic sensitivity assays were developed with a qualitative method based on diffusion (antibiotic-discs with 200 μM concentration of each antibiotic: ampicillin, gentamicin, methicilin and vancomycin) and a quantitative method based on microdilution (seven concentrations: 0, 3.12, 6.25, 12.5, 25, 50 and 100 μM of each antibiotic). Determination of the resistance (P-value < 0.05) was performed by a linear model comparing the growth of the clinical isolates against the reference strain.

Then, according to previous studies (Groot *et al*., 2012), microdilution assays were conducted to evaluate the antibacterial activity of the peptides following the guidelines suggested by the Clinical and Laboratory Standards Institute (CLSI, 2012). All the *S. aureus* mentioned above were exposed to the eight peptides individually and combined in the assay. Briefly, a standardized bacterial inoculum (OD 595 nm = 0.001) was exposed to seven different concentrations (0, 3.12, 6.25, 12.5, 25, 50 and 100 μM) of the peptides individually and combined in Luria Bertani (LB) Media (LB-Media, Miller, Sigma-Aldrich) in 96-well microplates for 6, 8 and 24 hours at 37ºC. The antibiotics ampicillin, gentamicin, methicilin and vancomycin were used as control and were tested in the same concentrations of the peptides. After incubation, the optic density was determined reading the microplates at 595nm in a spectrophotometer. Each peptide, the eight peptide combination and the control experiments were done with two replicates with three repetitions per concentration.

### Cell culture of fibroblasts (VERO) and human monocytes (THP-1) and differentiation to macrophages

The African green monkey fibroblast-like kidney cells (VERO) were maintained at 37 ºC and 5 % CO2 in DMEM media with 10 % FBS and 1 % Penicillin/Streptomycin. The human monocytic cell line (THP-1) was maintained at 2×10^5^ cells/mL in RMPI 1640 media with 10 % FBS and 1 % Penicillin/Streptomycin at 37 ºC and 5 % CO2. Macrophages were obtained using the protocol proposed by (Daigneault *et al*. 2010), exposing THP-1 monocytes to 200 nM phorbol 12-myristate 13-acetate (PMA, Sigma, P8139) for three days, followed by five days of incubation in RPMI media with supplements. After treatment, the confirmation of the monocytes-to-macrophages differentiation was performed by analysis of the expression of recognized macrophage markers (CD11b, CD14 and CD36). Briefly, cells were detached by exposing them to 10 mM EDTA for 40 minutes at 4 ºC. Supplemented media was added to the flasks and the cell suspension put in FACS tubes. Afterwards, two washes were performed by centrifuging for five minutes at 2500 rpm and 21 ºC and the pellet resuspended in 3 mL of PBS. The final suspension was incubated with 5 μL of APC Mouse Anti-Human CD11b (Clone D12, 333143, BD Biosciences), PE Mouse Anti-Human CD14 (Clone M5E2, 555398, BD Biosciences) and FITC Mouse Anti-Human CD36 (Clone CD38, 555454, BD Biosciences) for 30 minutes. After incubation, two more washes were performed and ultimately, the pellet with the marked cell was resuspended in 100 μL of PBS. Finally, cells were analyzed via flow cytometry (FACSCanto™, BD Biosciences).

### Determination of the cytotoxic activity of eight new peptides individually and combined in fibroblasts (VERO), monocytes (THP-1) and macrophages (THP-1 differentiated) cell cultures

MTT cytotoxic assays were realized with the three cell lines described above, including the differentiated macrophages. Firstly, 1×10^5^ cells/well were seeded in 96-well microplates for 24 hours. Following, they were exposed to seven concentrations (0, 3.12, 6.25, 12.5, 25, 50 and 100 μM) of the peptides, individually and combined, the antibiotics (ampicillin, gentamicin and vancomycin) and the positive control (doxorubicin). After incubation for 24 hours, MTT reagent (Sigma-Aldrich, St. Louis, USA) was added and the cells were incubated for two more hours. Then, the media was removed and 100 μL of DMSO were aggregated to each well and the plates were read at 595 nm with a spectrophotometer (Bio-Rad, Philadelphia, PA, USA). Ultimately, cytotoxic concentration 50 (CC50) was calculated as the concentration at which the peptides produced the 50 % of the death of the cells in two replicates with three repetitions each.

### Determination of the immunomodulatory activity of three new peptides individually in human macrophage (THP-1 differentiated) cell culture in three scenarios: without stimulation, with *E. coli* LPS stimulation and with *S. aureus* lysate stimulation

After differentiating the THP-1 monocytes to macrophages as mentioned previously, 1×10^5^ cells were seeded in 96-well plates. The macrophages were exposed to three concentrations (0, 6.25, 25 μM) of three peptides selected for their strong antibacterial and weak cytotoxic activity. Experiments were performed with and without 0.5 uL/mL of *Escherichia coli* LPS (Sigma-Aldrich, St. Louis, USA) or *S. aureus* lysate with two repetitions in two replicates. The *S. aureus* lysate was obtained previously by heat-killing (98 ºC for 30 minutes) the ON growth of the *S. aureus* reference strain (ATCC 25923), followed by the lyophilization of the pellet obtained after the centrifugation (2500 rpm for five minutes) of the bacterial culture and washing with ddH2O. Afterwards, the exposed macrophage culture plates were incubated for 48 hours at 37ºC with 5 % CO2 and the cells and supernatant were collected for further analysis. Firstly, in the supernatants, the concentration of 13 cytokines (IL-1β, IFN-α, IFN-γ, TNF-*α*, CCL2, IL-6, IL-8, IL-10, IL-12p70, IL-17A, IL-18, IL-23 e IL-33) was determined with the LEGENDplex™ Human Inflammation Panel kit 13-plex (Biolegend) following the recommendations of the kit. The results were obtained using flow cytometry (FACSCanto™, BD Biosciences). Secondly, the expression of the cell markers CD11b, CD14 and CD36 was evaluated in the macrophages following the methodology described above.

### Statistical Analysis

All statistical analysis were performed using the computing environment R (R Development Core Team, 2008). In all cases, a residual analysis was performed to evaluate the normality and homoscedasticity of the data and all plots were created with the package ggplot2 (Wickham, 2009). Also, for the evaluation of the antibacterial and cytotoxic activity, the GRmetric package (Clark *et al*., 2016 and Gentleman *et al*., 2004) was used for the creation of the dose-response sigmoidal models and calculations of the IC50, EC50 and CC50. In addition, the model selection analysis, Bartlett’s test and DFA analysis were performed using the packages MuMin (Gelman, 2008), psych (Revelle, 2017) and candisc (Friendly Fox, 2017) respectively.

## ACKNOWLEDGMENTS

We gratefully thank Camila Mejia from the Hematology Laboratory of Pontificia Universidad Javeriana for their support in the flow cytometry assays. We also thank Dr. Patricia Escobar and Dr. Laura Neira from the CINTROP of Universidad Industrial de Santander and Dr. Alejandro Hernandez from the GEMAPOL of Universidad del Quindío, for their kind donation of the THP-1 cell line. This research was supported by the “Proyecto Semilla” Convocatory of the Vicedecanatura de Investigaciones of the Sciences Faculty of Universidad de los Andes and the “Producto Terminado” Convocatory of the Vicerrectoría de Investigación of Universidad de los Andes.

## AUTHOR CONTRIBUTIONS

All the authors contributed to the design and implementation of the research, to the analysis of the results and to the writing of the manuscript.

---

**Supplementary Material Table 1.** Predicted stability features of the eight new peptides. The ProtParam Tool and PeptideCutter Tool of ExPASy and the HLP: “Web server for predicting half-life of the peptide in an intestine-like environment” were used to predict stability features of the eight new peptides. All the peptides were predicted to stay stable for hours *in vitro* in mammalian reticulocytes, while only AA, AT and GF might conserve their structure *in vivo* in yeast and bacteria. The instability index classified KS, GF, KV and LK as unstable in test tube conditions, while the aliphatic index predicted that RN might be the peptide with the highest potential to be thermostable. Interestingly, the predictions in an intestine-like environment showed, that all the peptides might be stable in a proteolytic environment. The PeptideCutter Tool predicted that NS would be susceptible to the Staphylococcal peptidase I.

| ID | Half-life<sup>1</sup> |  |  | Instability index<sup>1</sup> | Class<sup>1</sup> | Aliphatic index<sup>1</sup> | Intestin-like environment<sup>2</sup> |  | Cutters<sup>3</sup> |
| --- | --- | --- | --- | --- | --- | --- | --- | --- | --- |
|  | *in vitro* | *in vivo* |  |  |  |  | Half-life | Stability |  |
|  |  | Yeast | Bacteria |  |  |  |  |  |  |
| AA | 4.4h | >20h | >10h | -21,7 | Stable | 89,09 | 1,03 | High | LysC, LysN, Proteinase K, Thermolysin, Trypsin |
| KS | 1.3h | 3min | 3min | 119 | Unstable | 48,75 | 2,41 | High | CNBr, Chymotrypsin-low specificity, LysC, LysN, Pepsin (pH1.3 and >2), Proteinase K, Thermolysin, Trypsin |
| NS | 1.4h | 3min | >10h | 23,0 | Stable | 68,82 | 0,36 | Normal | Asp-N endopeptidase + N-terminal Glu, CNBr, Chymotrypsin-low specificity, Glutamyl endopeptidase, LysC, LysN, Pepsin (pH1.3 and >2), Proteinase K, Staphylococcal peptidase I, Thermolysin, Trypsin |
| RN | 1h | 2min | 2min | 12,4 | Stable | 148,5 | 0,74 | Normal | Arg-C proteinase, Chymotrypsin-low specificity, Clostripain, LysC, LysN, Pepsin (pH1.3 and >2), Proteinase K, Thermolysin, Trypsin |
| AT | 4.4h | >20h | >10h | -6,27 | Stable | 110,7 | 0,83 | Normal | CNBr, Chymotrypsin-high specificity, Chymotrypsin-low specificity, LysC, LysN, Pepsin (pH1.3 and >2), Proteinase K, Thermolysin, Trypsin |
| GF | 30h | >20h | >10h | 42,2 | Unstable | 106,3 | 0,52 | Normal | Arg-C proteinase, CNBr, Chymotrypsin-high specificity, Chymotrypsin-low specificity, Clostripain, LysC, LysN, Pepsin (pH1.3 and >2), Proteinase K, Thermolysin, Trypsin |
| KV | 1.3h | 3min | 3min | 44,9 | Unstable | 151 | 0,90 | Normal | Chymotrypsin-low specificity, LysC, Pepsin (pH1.3 and >2), Proteinase K, Thermolysin, Trypsin |
| LK | 5.5h | 3min | 2min | 52,4 | Unstable | 165,2 | 1,00 | Normal | Chymotrypsin-low specificity, LysC, LysN, Pepsin (pH1.3 and >2), Proteinase K, Thermolysin, Trypsin |

**Supplementary Material Table 2.**
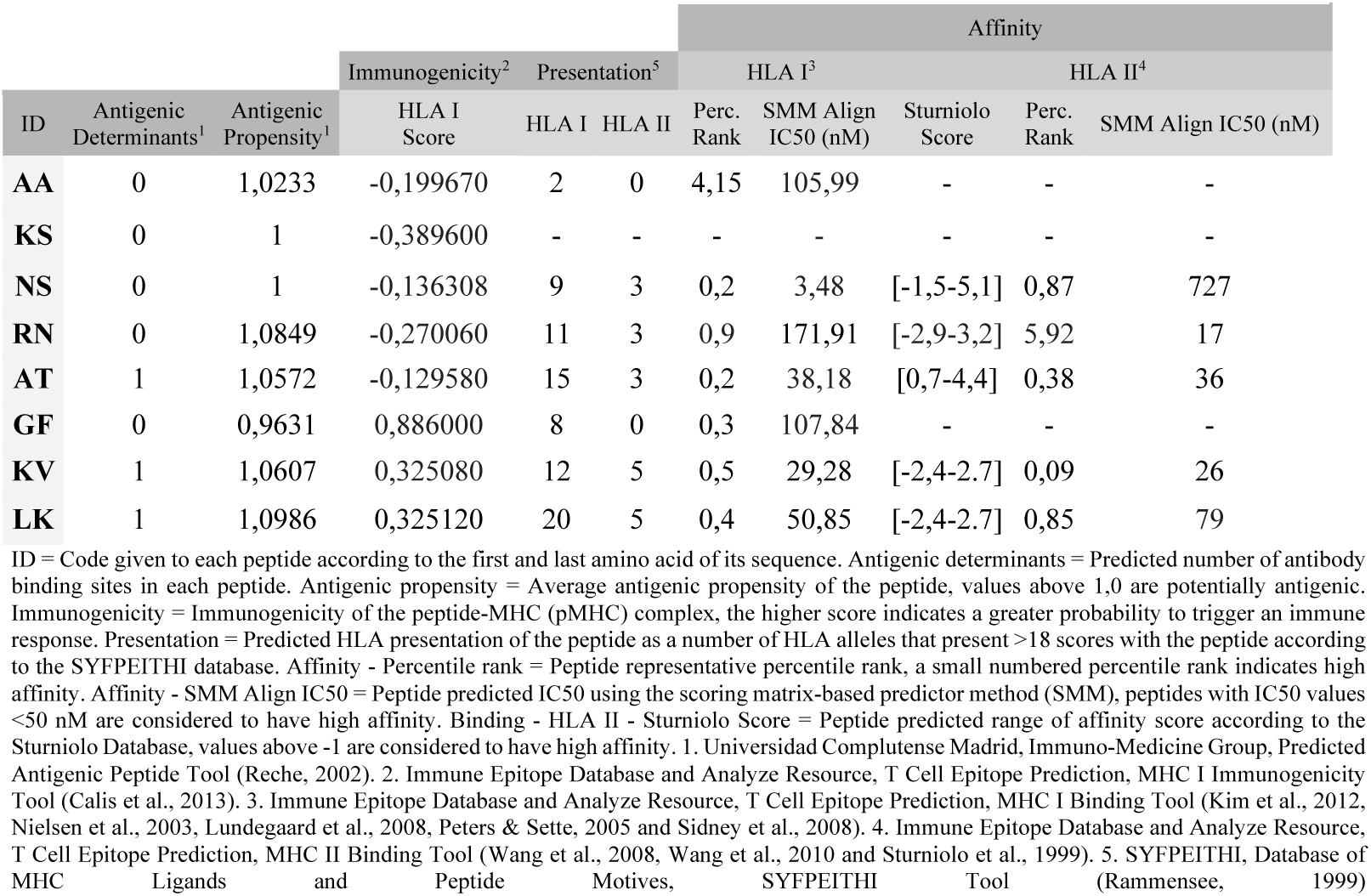
Antigenicity of eight new peptides. The Predicted Antigenic Peptide Tool, the Immune Epitope Database and Analyze Resource and the SYFPEITHI, Database of MHC Ligands and Peptide Motives were used to identify antigenicity features of the peptides considered as their potential to be presented or interact with the HLA (I and II). Seven of the peptides were classified as potentially antigenic, while only GF, KV, and LK were predicted to have antigenic determinants which might activate B cells. Results about the immunogenicity ranked these last three peptides also as the peptides with a higher potential to trigger an immune response via HLA I. In addition, RN, AT, KV and LK might be presented by >10 HLA I alleles, while KV and LK would be presented by >4 HLA II alleles. In regards to the peptide affinity to HLA molecules, NS, AT, KV and LK peptides might bind with high affinity to HLA type I and RN, AT, KV and LK to the HLA type II. Results of potential presentation and affinity of the KS peptides to HLA (I and II) molecules were not developed because the peptide sequence was too short. This was also the case for the AA and GF analysis prediction for their affinity to HLA (I and II) molecules.

**Supplementary Material Table 3.**
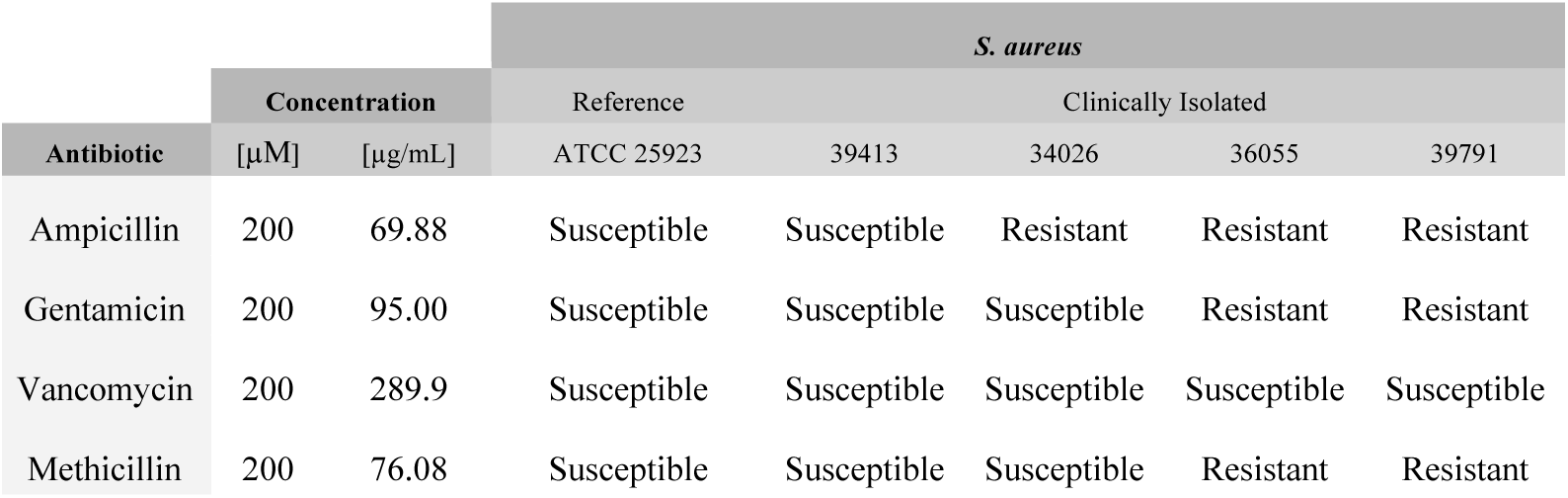
Qualitative classification of the resistance profile of each of the *S. aureus* used in this study to the four antibiotics tested. Antibiotic sensitivity assays were performed using a qualitative method based on diffusion, antibiotic-discs with 200μM concentration of each antibiotic were seeded in LB agar plates where the bacteria was grown. The concentration used is shown in μM and μg/mL units for each antibiotic. Results resembled what was expected and the quantitative results are shown below.

**Supplementary Material Table 4.** Quantitative results of the resistance profile of each of the *S. aureus* used in this study to the four antibiotics tested. Microdilution assays were performed exposing the bacteria to the antibiotics at concentrations between 3.12 and 100 μM, results were obtained after 6, 8 and 24 hours of exposure. The half maximal inhibitory concentration (IC50), and the half maximal effective concentration (EC50) are shown. In all cases the significance was <0.05 and the coefficient of determination of the model (r2) > 0.77.

| *S. aureus* | Hours of exposure | Antibiotic |  |  |  |  |  |  |  |
| --- | --- | --- | --- | --- | --- | --- | --- | --- | --- |
|  |  | Ampicillin |  | Gentamicin |  | Methicillin |  | Vancomycin |  |
|  |  | IC50 (uM) | EC50 (uM) | IC50 (uM) | EC50 (uM) | IC50 (uM) | EC50 (uM) | IC50 (uM) | EC50 (uM) |
| ATCC 25923 | 6 | 2,24 | 2,06 | 1,27 | 1,23 | 0.566 | 0.551 | 0,98 | 0,96 |
|  | 8 | 1,68 | 1,6 | 1,12 | 1,09 | 0,299 | 0,295 | 1,05 | 1,04 |
|  | 24 | 1,22 | 1,17 | 1,27 | 1,23 | 0.521 | 0.515 | 0,9 | 0,89 |
| Clinically isolated 39413 | 6 | 3,88 | 3,74 | 1,59 | 1,51 | 0.322 | 0.318 | 1,56 | 1,56 |
|  | 8 | 2,98 | 2,98 | 1,06 | 1,04 | 0,640 | 0,635 | 0,95 | 0,94 |
|  | 24 | 8,73 | 8,74 | 4,58 | 4,58 | 0.419 | 0.416 | 0,62 | 0,61 |
| Clinically isolated 34026 | 6 | 17,47 | 17,5 | 1,57 | 1,5 | 2.15 | 2.13 | 1,13 | 1,11 |
|  | 8 | 26,51 | 26,5 | 1,06 | 1,01 | 1,52 | 1,50 | 0,85 | 0,84 |
|  | 24 | 32,67 | 32,7 | 12,3 | 12,3 | 9.66 | 8.49 | 1,08 | 1,08 |
| Clinically isolated 36055 | 6 | 125 | 125 | - | 66,1 | - | 2.09 | 0,92 | 0,92 |
|  | 8 | 42,1 | 24,2 | - | - | - | - | 0,7 | 0,69 |
|  | 24 | - | 27,9 | - | - | - | 0.26 | 0,97 | 0,96 |
| Clinically isolated 39791 | 6 | 58,5 | 32,1 | - | 18,5 | - | 21.62 | 1,07 | 1,07 |
|  | 8 | 32,94 | 9,56 | - | - | - | - | 0,82 | 0,81 |
|  | 24 | - | 12,76 | - | 15,3 | - | 0.247 | 0,88 | 0,88 |

**Supplementary Material Table 5.**
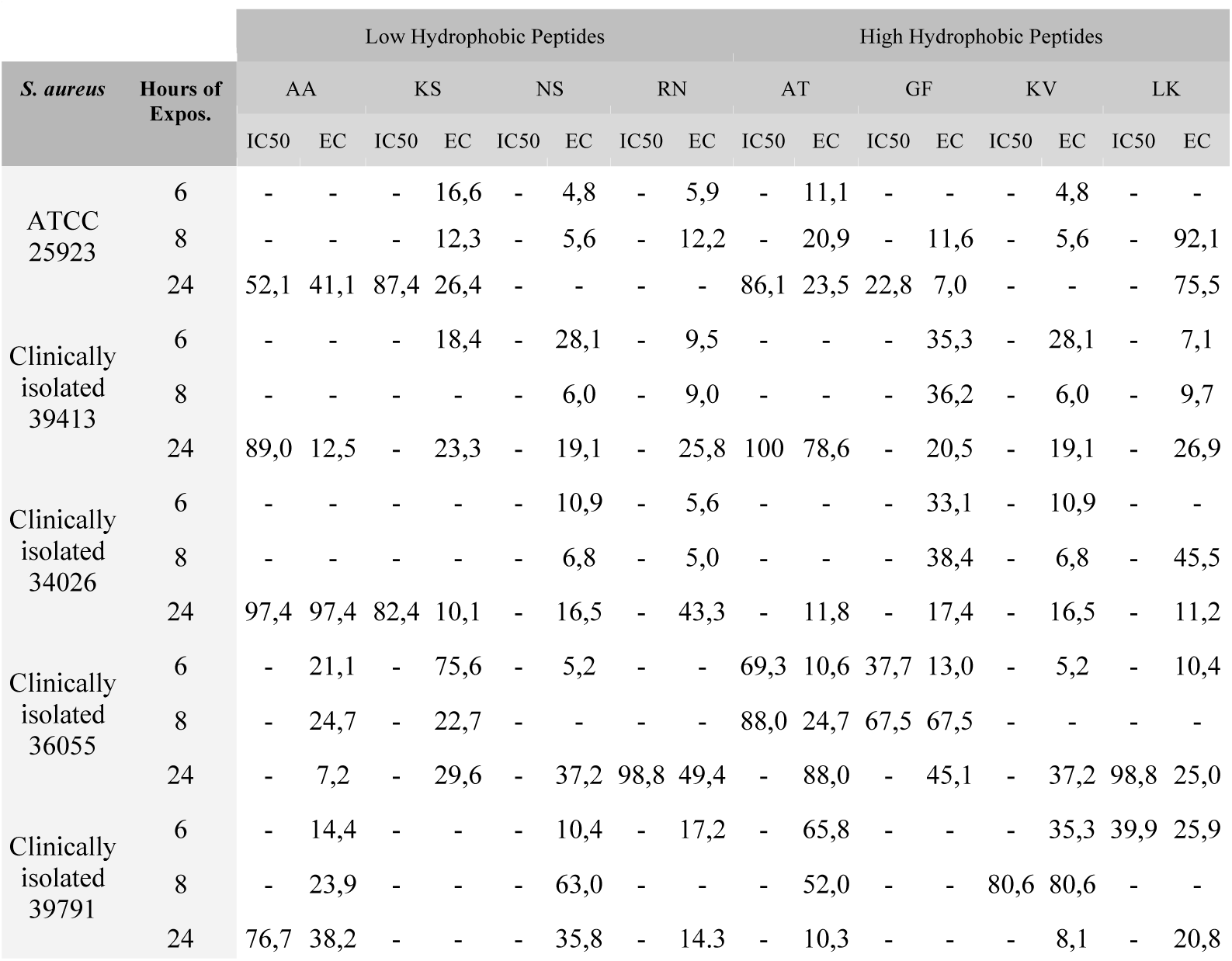
Antimicrobial activity of eight new peptides individually. Microdilution assays were performed exposing the *S. aureus* to the peptides individually at concentrations between 3.12 and 100 μM, results were obtained after 6, 8 and 24 hours of exposure. In all cases the significance was <0.05 and the coefficient of determination of the model (r2) > 0.77.

**Supplementary Material Table 6.**
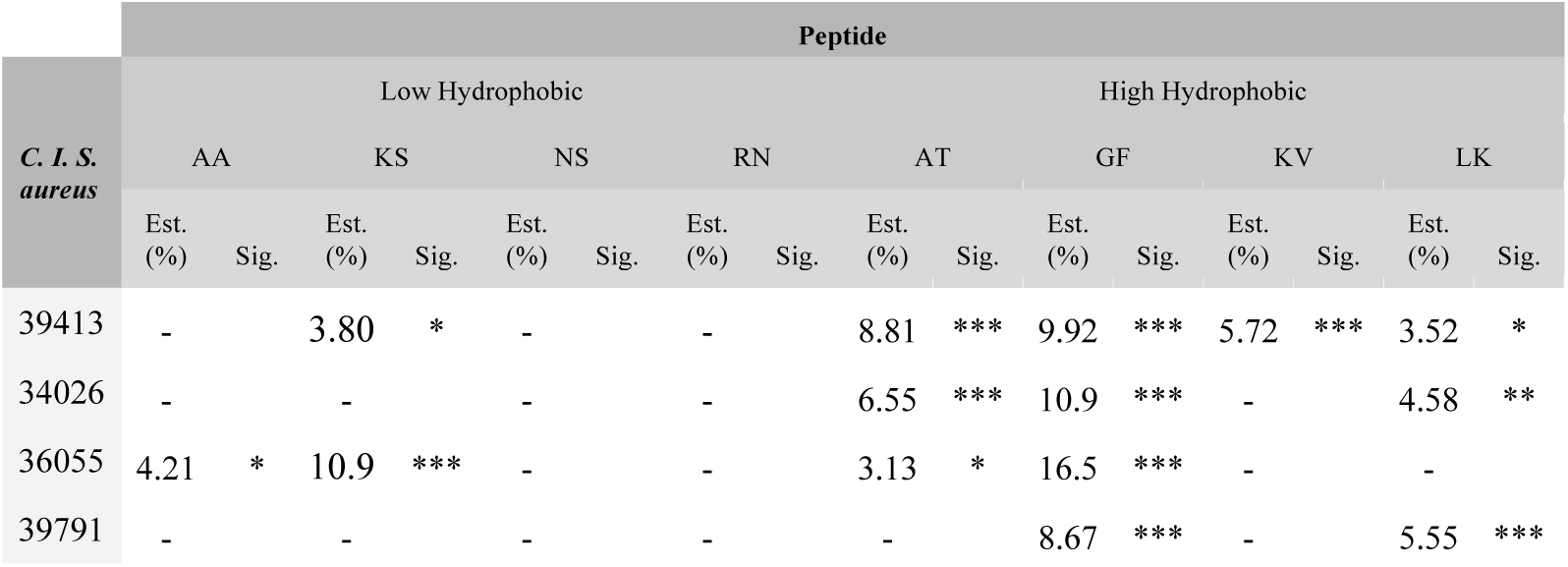
Comparison of the antimicrobial activity of eight new peptides individually between the clinically isolated *S. aureus*. The antibacterial activity of the peptides individually against the four clinically isolated were compared with the corresponding antibacterial activity identified for the reference *S. aureus* strain. Only results after 24 hours of exposure are shown. The estimate (decrease (%) in the antibacterial activity) and the significance (P-value = ***<0.001, **<0.01, *<0.05) are shown. In all cases, the coefficient of determination of the model (r2) was >0.57.

**Supplementary Material Table 7.** Comparison of the antimicrobial activity of eight new peptides individually with antibiotics. Microdilution assays were performed exposing the *S. aureus* to the peptides individually at concentrations between 3.12 and 100μM, results were obtained after 6, 8 and 24 hours of exposure. The half maximal inhibitory concentration (IC50), and the half maximal effective concentration (EC50) are shown. In all cases the significance was <0.05 and the coefficient of determination of the model (r2) > 0.77.

| *S. aureus* | Antibiotic | Peptide |  |  |  |  |  |  |  |  |  |  |  |  |  |  |  |
| --- | --- | --- | --- | --- | --- | --- | --- | --- | --- | --- | --- | --- | --- | --- | --- | --- | --- |
|  |  | Low Hydrophobic |  |  |  |  |  |  |  | High Hydrophobic |  |  |  |  |  |  |  |
|  |  | AA |  | KS |  | NS |  | RN |  | AT |  | GF |  | KV |  | LK |  |
|  |  | Est. | Sig. | Est. | Sig. | Est. | Sig. | Est. | Sig. | Est. | Sig. | Est. | Sig. | Est. | Sig. | Est. | Sig. |
| ATCC 25923 | Ampicillin | -46.4 | *** | -49.0 | *** | -52.6 | *** | -53.8 | *** | -45.6 | *** | -36.6 | *** | -50.5 | *** | -48.9 | *** |
|  | Gentamicin | -45.5 | *** | -48.2 | *** | -51.8 | *** | -52.9 | *** | -44.8 | *** | -35.8 | *** | -49.7 | *** | -48.1 | *** |
|  | Methicillin | -49.7 | *** | -52.3 | *** | -55.9 | *** | -57.1 | *** | -48.9 | *** | -39.9 | *** | -53.8 | *** | -52.2 | *** |
|  | Vancomycin | -50.4 | *** | -53.1 | *** | -56.7 | *** | -57.8 | *** | -49.7 | *** | -40.6 | *** | -54.6 | *** | -52.9 | *** |
| C. I. 39413 | Ampicillin | -26.5 | *** | -32.5 | *** | -27.2 | *** | -28.2 | *** | -34.2 | *** | -25.1 | *** | -35.9 | *** | -32.1 | *** |
|  | Gentamicin | -38.2 | *** | -44.2 | *** | -38.8 | *** | -39.8 | *** | -45.8 | *** | -36.8 | *** | -47.6 | *** | -43.8 | *** |
|  | Methicillin | -51.0 | *** | -57.0 | *** | -51.7 | *** | -52.6 | *** | -58.6 | *** | -49.6 | *** | -60.4 | *** | -56.6 | *** |
|  | Vancomycin | -51.3 | *** | -57.3 | *** | -51.9 | *** | -52.9 | *** | -58.9 | *** | -49.9 | *** | -60.7 | *** | -56.9 | *** |
| C. I. 34026 | Ampicillin | - |  | - | *** | -12.0 | *** | - | *** | -11.9 | * | - |  | -12.9 | * | -13.2 | * |
|  | Gentamicin | -24.9 | *** | -18.7 | *** | -27.8 | *** | -24.9 | *** | -27.8 | *** | -23.1 | *** | -28.8 | *** | -29.1 | *** |
|  | Methicillin | -24.7 | *** | -18.5 | *** | -27.6 | *** | -24.7 | *** | -27.5 | *** | -22.8 | *** | -28.5 | *** | -28.8 | *** |
|  | Vancomycin | -53.4 | *** | -47.2 | *** | -56.3 | *** | -53.3 | *** | -56.2 | *** | -51.5 | *** | -57.2 | *** | -57.5 | *** |
| C. I. 36055 | Ampicillin | - |  | - |  | -14.7 | *** | - |  | - |  | - |  | - |  | - |  |
|  | Gentamicin | 12.1 | ** | - |  | -9.90 | *** | 13.3 | ** | 13.5 | ** | 9.66 | * | 10.9 | * | 12.7 | ** |
|  | Methicillin | -10.7 | * | -20.1 | *** | -13.0 | *** | -9.60 | *** | -9.35 | ** | -13.2 | ** | -11.9 | ** | -10.1 | * |
|  | Vancomycin | -54.2 | *** | -63.6 | *** | -56.4 | *** | -53.0 | *** | -52.7 | *** | -56.6 | *** | -55.4 | *** | -53.5 | *** |
| C. I. 39791 | Ampicillin | - |  | - |  | - |  | - |  | - |  | - |  | - |  | - |  |
|  | Gentamicin | - |  | - |  | - |  | - |  | - |  | - |  | - |  | - |  |
|  | Methicillin | - |  | - |  | -9.78 | * | -8.57 | * | - |  | - |  | - |  | -11.6 | ** |
|  | Vancomycin | -50.2 | *** | -55.1 | *** | -56.8 | *** | -55.5 | *** | -52.8 | *** | -49.3 | *** | -53.9 | *** | -58.5 | *** |

**Supplementary Material Table 8.** Antibacterial activity of eight new peptides combined. Microdilution assays were performed exposing the *S. aureus* to the peptides in combination at concentrations between 3.12 and 100 μM, results were obtained after 6, 8 and 24 hours of exposure. The antibacterial activity of the peptide combination was compared with the antibiotic controls. The half maximal inhibitory concentration (IC50), the half maximal effective concentration (EC50), the estimate (Estim.) (difference shown compared to the antibiotic (%)) and the significance (Sign.) (P-value = ***<0.001, **<0.01, *<0.05) are shown.

| *S. aureus*<br>Strain | Hours of<br>exposure | Eight<br>Peptides<br>combined |  | Comparison results with antibiotics |  |  |  |  |  |  |  |
| --- | --- | --- | --- | --- | --- | --- | --- | --- | --- | --- | --- |
|  |  |  |  | Ampicillin |  | Gentamycin |  | Methicillin |  | Vancomycin |  |
|  |  | IC50 | EC50 | Estim. | Sign. | Estim. | Sign. | Estim. | Sign. | Estim. | Sign. |
| ATCC<br>25923 | 6 | 58,6 | 8,4 | -33.4 | *** | -41.0 | *** | -42.6 | *** | -43.7 | *** |
|  | 8 | 52,4 | 52,4 | -41.9 | *** | -46.6 | *** | -48.8 | *** | -48.3 | *** |
|  | 24 | 33,3 | 33,3 | -40.8 | *** | -39.9 | *** | -44.1 | *** | -44.8 | *** |
| Clinically<br>isolated<br>39413 | 6 | - | 12,3 | -34.4 | *** | -43.8 | *** | -50.3 | *** | -47.2 | *** |
|  | 8 | - | 5,6 | -33.9 | *** | -42.7 | *** | -45.8 | *** | -44.7 | *** |
|  | 24 | - | 11,1 | -24.3 | *** | -35.9 | *** | -48.7 | *** | -49.0 | *** |
| Clinically<br>isolated<br>34026 | 6 | - | 17,1 | -13.1 | * | -41.6 | *** | -45.6 | *** | -45.8 | *** |
|  | 8 | 58,2 | 6,8 | - |  | -40.9 | *** | -41.0 | *** | -43.1 | *** |
|  | 24 | - | 3,7 | - |  | -19.4 | *** | -19.2 | *** | -47.9 | *** |
| Clinically<br>isolated<br>36055 | 6 | 15,8 | 2,1 | **23.6** | *** | **21.1** | *** | **8.80** | * | -37.0 | *** |
|  | 8 | 7,7 | 2,0 | **11.9** | ** | **32.4** | *** | **20.4** | *** | -32.6 | *** |
|  | 24 | 90,8 | 5,5 | **17.2** | *** | **22.4** | *** | - |  | -43.9 | *** |
| Clinically<br>isolated<br>39791 | 6 | 14,0 | 3,8 | **11.3** | ** | **18.1** | *** | **11.2** | ** | -34.8 | *** |
|  | 8 | 9,3 | 2,4 | - |  | **29.1** | *** | **25.9** | *** | -32.7 | *** |
|  | 24 | 56,7 | 3,2 | **10.2** | * | **14.9** | *** | - |  | -42.6 | *** |

**Supplementary Material Table 9.** Cytotoxic activity of eight new peptides individually and combined. MTT assays were performed, exposing three cell lines to the peptides individually and combined at concentrations between 3.12 and 100μM. Results were obtained after 24 hours of exposure. The half maximal cytotoxic concentration (CC50) is shown. In all cases the significance was <0.05 and the coefficient of determination of the model (r2) > 0.50.

| Cell Line | Low Hydrophobic Peptides |  |  |  | High Hydrophobic Peptides |  |  |  | Eight Peptide Combined | Doxorubicin |
| --- | --- | --- | --- | --- | --- | --- | --- | --- | --- | --- |
|  | AA | KS | NS | RN | AT | GF | KV | LK |  |  |
| | CC50 (μM) | CC50 (μM) | CC50 (μM) | CC50 (μM) | CC50 (μM) | CC50 (μM) | CC50 (μM) | CC50 (μM) | CC50 (μM) | CC50 (μM) |
| VERO fibroblasts | >100 | >100 | >100 | >100 | >100 | >100 | >100 | >100 | >100 | 56.7 |
| THP-1 monocytes | >100 | >100 | >100 | >100 | 63.5 | >100 | >100 | 51.6 | 21.8 | 5.64 |
| THP-1 macrophages | >100 | >100 | >100 | >100 | >100 | >100 | >100 | >100 | >100 | 5.26 |

**Supplementary Material Table 10.** Immunomodulatory activity of three new peptides. After exposing the peptides to the macrophages in three different stimulation scenarios and at different concentrations (6.25 and 25μ M), the cytokine concentration in the cell culture media was calculated. Results were obtained after 48 hours of exposure. The cytokine concentration, as well as the standard deviation are shown. Furthermore, the significance of the difference between the peptide concentrations are shown (P-value = ***<0.001, **<0.01, *<0.05).

| Peptide | Stimulation Scenario | Cytokine |  |  |  |  |  |  |  |  |  |  |  |
| --- | --- | --- | --- | --- | --- | --- | --- | --- | --- | --- | --- | --- | --- |
| | | IL-1β | | | CCL-2 | | | TNF-α | | | IL-8 | | |
|  |  | Without stimulation | *E. coli* LPS | *S. aureus* lysate | Without stimulation | *E. coli* LPS | *S. aureus* lysate | Without stimulation | *E. coli* LPS | *S. aureus* lysate | Without stimulation | *E. coli* LPS | *S. aureus* lysate |
| Without Peptide | Cytokine [pg/mL] | 1195.9 | 1853.4 | 1128.1 | 9109.6 | 8058.4 | 9052.5 | 640.8 | 1188.2 | 453.2 | 8058.3 | 8057.7 | 8078.9 |
|  | *S. D.* (pg/mL) | 100.4 | 127.5 | 4.9 | 0.985 | 5.67 | 5.18 | 24.5 | 18.9 | 22.0 | 9.99 | 2.96 | 5.24 |
| AA [6.25 μM] | Cytokine [pg/mL] | **2031.4** | **1442.0** | **1811.9** | 9048.7 | 9075.5 | 9098.8 | **922.7** | **541.9** | **903.0** | 8063.3 | 8036.0 | **8043.0** |
|  | *S. D.* (pg/mL) | **46.0** | **7.29** | **46.5** | 3.75 | 3.87 | 7.67 | **51.0** | **10.4** | **21.5** | 2.86 | 1.09 | **6.26** |
| AA [25 μM] | Cytokine [pg/mL] | **2170.2** | **1986.5** | **1770** | 9046.4 | 9047.5 | 9080.4 | **813.5** | **661.9** | **1028.9** | 8069.3 | 8066.4 | **8105.8** |
|  | *S. D.* (pg/mL) | **7.34** | **22.4** | **28.6** | 2.60 | 1.06 | 7.88 | **51.6** | **2.64** | **26.8** | 3.32 | 5.35 | **8.85** |
|  | Significance | ** | *** | * |  |  |  | *** | *** | *** |  |  | ** |
| GF [6.25 μM] | Cytokine [pg/mL] | **2098.8** | **1540.9** | **1799.8** | 9085.6 | **9074.9** | **9073.3** | **1028.4** | **678.5** | **725.9** | 8110.3 | 8061.5 | **8069.7** |
|  | *S. D.* (pg/mL) | **55.8** | **25.9** | **16.5** | 3.57 | **5.67** | **9.69** | **72.5** | **1.59** | **13.9** | 3.20 | 3.34 | **3.83** |
| GF [25 μM] | Cytokine [pg/mL] | **3146.9** | **2329.2** | **3201.3** | 9087.1 | **9053.0** | **9081.8** | **1507.7** | **784.6** | **1390.9** | 8120.5 | 8107.2 | **8303.6** |
|  | *S. D.* (pg/mL) | **122.9** | **132.5** | **120.1** | 6.59 | **4.38** | **1.19** | **167.5** | **33.6** | **86.1** | 10.3 | 6.74 | **44.6** |
|  | Significance | *** | *** | *** |  | * | * | *** | *** | *** |  |  | *** |
| RN [6.25 μM] | Cytokine [pg/mL] | **2876.4** | **1605.1** | **2113.0** | **9158.4** | **9116.5** | **9069.2** | **1729.7** | **683.8** | **739.2** | **8116.9** | **8038.3** | 8069.8 |
|  | *S. D.* (pg/mL) | **10.3** | **4.47** | **149.3** | **6.15** | **4.30** | **5.77** | **14.9** | **5.01** | **5.95** | **13.3** | **4.54** | 4.94 |
| RN [25 μM] | Cytokine [pg/mL] | **1447.0** | **3255.0** | **2658.1** | **9055.1** | **9165.5** | **9080.9** | **633.8** | **1935.5** | **1099.7** | **8038.4** | **8188.5** | 8071.9 |
|  | *S. D.* (pg/mL) | **72.1** | **110.3** | **23.4** | **8.12** | **14.2** | **3.29** | **16.3** | **235.6** | **13.5** | **1.27** | **16.1** | 726 |
|  | Significance | *** | *** | *** | *** | *** | * | *** | *** | *** | *** | *** |  |

## Bibliography

Apponyi, M. A., Pukala, T. L., Brinkworth, C. S., Maselli, V. M., Bowie, J. H., Tyler, M.J. Doyle, J. (2004). Host-defence peptides of Australian anurans: structure, mechanism of action and evolutionary significance. Peptides, 25(6), 1035–1054.

Attoub, S., Mechkarska, M., Sonnevend, A., Radosavljevic, G., Jovanovic, I., Lukic, M. L. Conlon, J. M. (2013). Esculentin-2CHa: a host-defense peptide with differential cytotoxicity against bacteria, erythrocytes and tumor cells. Peptides, 39, 95–102.

Badosa, E., Ferre, R., Planas, M., Feliu, L., Besalú, E., Cabrefiga, J. Montesinos, E. (2007). A library of linear undecapeptides with bactericidal activity against phytopathogenic bacteria. Peptides, 28(12), 2276–2285.

Bamberger, M.D. Boyd, S.E. (2005). Management of *Staphylococcus aureus* Infections. American Family Physician, 72(12), 2474–2481.

Brown, A. F., Leech, J. M., Rogers, T. R. McLoughlin, R. M. (2014). *Staphylococcus aureus* colonization: modulation of host immune response and impact on human vaccine design. Frontiers in Immunology, 4, 507.

Calis, J.J.A., Maybeno, M., Greenbaum, J.A., Weiskopf, D., De Silva, A.D., Sette, A., Kesmir, C., Peters, B. (2013). Properties of MHC class I presented peptides that enhance immunogenicity. PloS Computational Biology. 8(1):361.

Čeřovský, V., Buděšínský, M., Hovorka, O., Cvačka, J., Voburka, Z., Slaninová, J. Straka, J. (2009). Lasioglossins: three novel antimicrobial peptides from the venom of the eusocial bee *Lasioglossum laticeps* (Hymenoptera: Halictidae). ChemBioChem, 10(12), 2089–2099.

Chen, C., Li, Z., Huang, H., Suzek, B. E., Wu, C. H., UniProt Consortium. (2013). A fast peptide match service for UniProt knowledgebase. Bioinformatics, btt484.

Clark, N., Hafner, M., Kouril, M., Muhlich, J., Niepel, M., Williams, E., Sorger, P. Medvedovic, M. (2016). GRcalculator: an online tool for calculating and mining drug response data. doi: 10.6084/m9.figshare.4244408.v1

CLSI. (2012). M07-A9: Methods for dilution antimicrobial susceptibility tests for bacteria that grow aerobically. Clinical and Laboratory Standards Institute, 32(2).

Conlon, J. M. (2011). The contribution of skin antimicrobial peptides to the system of innate immunity in anurans. Cell and Tissue Research, 343(1), 201–212.

Conlon, J. M., Demandt, A., Nielsen, P. F., Leprince, J., Vaudry, H., Woodhams, D. C. (2009). The alyteserins: two families of antimicrobial peptides from the skin secretions of the midwife toad *Alytes obstetricans* (Alytidae). Peptides, 30(6), 1069–1073.

Conlon, J. M., Mechkarska, M., Pantic, J. M., Lukic, M. L., Coquet, L., Leprince, J., Nielsen, P. F. Rinaldi, A. C. (2013). An immunomodulatory peptide related to frenatin 2 from skin secretions of the Tyrrhenian painted frog *Discoglossus sardus* (Alytidae). Peptides, 40, 65–71.

Conlon, J. M., Mechkarska, M., Radosavljevic, G., Attoub, S., King, J. D., Lukic, M. L. McClean, S. (2014). A family of antimicrobial and immunomodulatory peptides related to the frenatins from skin secretions of the Orinoco lime frog *Sphaenorhynchus lacteus* (Hylidae). Peptides, 56, 132–140.

Daigneault, M., Preston, J. A., Marriott, H. M., Whyte, M. K. Dockrell, D. H. (2010). The identification of markers of macrophage differentiation in PMA-stimulated THP-1 cells and monocyte-derived macrophages. PloS One, 5(1), e8668.

Daley, M., Williams, T., Coyle, P., Furda, G., Dougherty, R. Hayes, P. (1993). Prevention and treatment of *Staphylococcus aureus* infections with recombinant cytokines. Cytokine, 5(3), 276–284.

Friendly, M. Fox, J. (2017). Candisc: Visualizing Generalized Canonical Discriminant and Canonical Correlation Analysis

Fournier, B. Philpott, D. J. (2005). Recognition of *Staphylococcus aureus* by the innate immune system. Clinical Microbiology Reviews, 18(3), 521–540.

Fox, J. L. (2013). Antimicrobial peptides stage a comeback. Nature News Feature.

Gasteiger, E., Hoogland, C., Gattiker, A., Duvaud, S., Wilkins, M. R., Appel, R. D. Bairoch, A. (2005). Protein Identification and Analysis Tools on the ExPASy Server. The Proteomics Protocols Handbook, Humana Press, 571–607

Gentleman, R. C., Carey, V. J., Bates, D. M., Bolstad, B., Dettling, M., Dudoit, S., Ellis, B., Gautier, L., Ge, Y., Gentry, J., Hornik, K., Hothorn, T., Huber, W., Iacus, S., Irizarry, R., Leisch, F., Li, C., Maechler, M., Rossini, A. J., Sawitzki, G., Smith, C., Smyth, G., Tierney, L., Yang, J. Y. Zhang, J. (2004). Bioconductor: open software development for computational biology and bioinformatics. Genome Biology, 5, R80.

Gelman, A. (2008) Scaling regression inputs by dividing by two standard deviations. Statistics in medicine, 27, 2865–2873.

Groot, H., Muñoz-Camargo, C., Moscoso, J., Riveros, G., Salazar, V., Florez, F. K. Mitrani, E. (2012). Skin micro-organs from several frog species secrete a repertoire of powerful antimicrobials in culture. The Journal of Antibiotics, 65(9), 461–467.

Guaní-Guerra, E., Santos-Mendoza, T., Lugo-Reyes, S. O. Terán, L. M. (2010). Antimicrobial peptides: general overview and clinical implications in human health and disease. Clinical Immunology, 135(1), 1–11.

Hancock, R. E., Nijnik, A. Philpott, D. J. (2012). Modulating immunity as a therapy for bacterial infections. Nature Reviews Microbiology, 10(4), 243–254.

Khamis, A. M., Essack, M., Gao, X. Bajic, V. B. (2014). Distinct profiling of antimicrobial peptide families. Bioinformatics, 31(6), 849–856.

Kim, H. S., Yoon, H., Minn, I., Park, C. B., Lee, W. T., Zasloff, M. Kim, S. C. (2000). Pepsin-mediated processing of the cytoplasmic histone H2A to strong antimicrobial peptide Buforin I. The Journal of Immunology, 165(6), 3268–3274.

Kim, Y., Sidney, J., Pinilla, C., Sette, A. Peters, B. (2009). Derivation of an amino acid similarity matrix for peptide: MHC binding and its application as a Bayesian prior. BMC Bioinformatics, 1(394).

Kluytmans, J., Van Belkum, A. Verbrugh, H. (1997). Nasal carriage of *Staphylococcus aureus*: epidemiology, underlying mechanisms, and associated risks. Clinical Microbiology Reviews, 10(3), 505–520.

Kozlov, S. A., Vassilevski, A. A., Feofanov, A. V., Surovoy, A. Y., Karpunin, D. V., Grishin, E. V. (2006). Latarcins, antimicrobial and cytolytic peptides from the venom of the spider *Lachesana tarabaevi* (Zodariidae) that exemplify biomolecular diversity. Journal of Biological Chemistry, 281(30), 20983–20992.

Lai, R., Zheng, Y. T., Shen, J. H., Liu, G. J., Liu, H., Lee, W. H. Zhang, Y. (2002). Antimicrobial peptides from skin secretions of Chinese red belly toad *Bombina maxima*. Peptides, 23(3), 427–435.

Liu, C., Bayer, A., Cosgrove, S. E., Daum, R. S., Fridkin, S. K., Gorwitz, R. J., Kaplan, S. L., Karchmer, A. W., Levine, D. P., Murray, B. E., Rybak, M. J., Talan, D. A. Chambers, H. F. (2011). Clinical practice guidelines by the Infectious Diseases Society of America for the treatment of methicillin-resistant *Staphylococcus aureus* infections in adults and children. Clinical Infectious Diseases, 52(3), e18–e55.

Liu, H., Perlman, H., Pagliari, L. J. Pope, R. M. (2001). Constitutively activated Akt-1 is vital for the survival of human monocyte-differentiated macrophages. Journal of Experimental Medicine, 194(2), 113–126.

Lundegaard, C., Lamberth, K., Harndahl, M., Buus, S., Lund, O. Nielsen, M. (2008). NetMHC-3.0: Accurate web accessible predictions of Human, Mouse, and Monkey MHC class I affinities for peptides of length 8-11. NAR, 36, 509–512.

Miller, L. S., Pietras, E. M., Uricchio, L. H., Hirano, K., Rao, S., Lin, H., O’Connell, R. M., Iwakura, Y., Cheung, A. L., Cheng, G. Modlin, R. L. (2007). Inflammasome-mediated production of IL-1β is required for neutrophil recruitment against *Staphylococcus aureus In vivo*. The Journal of Immunology, 179(10), 6933–6942.

Meng, H. Kumar, K. (2007). Antimicrobial activity and protease stability of peptides containing fluorinated amino acids. Journal of the American Chemical Society, 129(50), 15615–15622.

Mohamed, M. F., Abdelkhalek, A. Seleem, M. N. (2016). Evaluation of short synthetic antimicrobial peptides for treatment of drug-resistant and intracellular *Staphylococcus aureus*. Scientific reports, 6, 29707.

Montgomery, C. P., Daniels, M. D., Zhao, F., Spellberg, B., Chong, A. S. Daum, R. S. (2013). Local inflammation exacerbates the severity of *Staphylococcus aureus* skin infection. PloS One, 8(7), e69508.

Muñoz-Camargo, C., Méndez, M. C., Salazar, V., Moscoso, J., Narváez, D., Torres, M. M., Florez, F. K., Groot, H. Mitrani, E. (2016). Frog skin cultures secrete anti-yellow fever compounds. The Journal of Antibiotics.

Myles, I. A., Fontecilla, N. M., Valdez, P. A., Vithayathil, P. J., Naik, S., Belkaid, Y., Ouyang, W. Datta, S. K. (2013). Signaling via the IL-20 receptor inhibits cutaneous production of IL-1β and IL-17A to promote infection with methicillin-resistant *Staphylococcus aureus*. Nature Immunology, 14(8), 804–811.

Nguyen, L. T., Haney, E. F. Vogel, H. J. (2011). The expanding scope of antimicrobial peptide structures and their modes of action. Trends in biotechnology, 29(9), 464–472.

Nielsen, M., Lundegaard, C., Worning, P., Lauemøller, S. L., Lamberth, K., Buus, S., Brunak, S. Lund, O. (2003). Reliable prediction of T-cell epitopes using neural networks with novel sequence representations. Protein Science, 12, 1007–1017.

Park, C. B., Kim, M. S. Kim, S. C. (1996). A Novel Antimicrobial Peptide from *Bufo gargarizans*. Biochemical and Biophysical Research Communications, 218(1), 408–413.

Patel, J. B., Gorwitz, R. J. Jernigan, J. A. (2009). Mupirocin resistance. Clinical Infectious Diseases, 49(6), 935–941.

Peters, B. Sette, A. (2005). Generating quantitative models describing the sequence specificity of biological processes with the stabilized matrix method. BMC Bioinformatics, 6, 132.

R Development Core Team. (2008). R: A language and environment for statistical computing. R Foundation for Statistical Computing, Vienna, Austria. ISBN 3-900051-07-0.

Rammensee, H. G., Bachmann, J., Emmerich, N. P. N., Bachor, O. A., Stevanović, S. (1999). SYFPEITHI: database for MHC ligands and peptide motifs. Immunogenetics, 50(3-4), 213–219.

Reche, P. A. (2002). Predicted Antigenic Peptides Tool. Immuno-Medicine Group. Universidad Complutense Madrid

Revelle, W. (2017). Psych: procedures for personality and psychological research. Northwestern University, Evanston, Illinois, USA, https://CRAN.R-project.org/package=psych

Roelants, K., Fry, B. G., Ye, L., Stijlemans, B., Brys, L., Kok, P. Bossuyt, F. (2013). Origin and functional diversification of an amphibian defense peptide arsenal. PLoS Genetics, 9(8), e1003662.

Rollins-Smith, L. A., Carey, C., Longcore, J., Doersam, J. K., Boutte, A., Bruzgal, J. E. Conlon, J. M. (2002). Activity of antimicrobial skin peptides from ranid frogs against *Batrachochytrium dendrobatidis*, the chytrid fungus associated with global amphibian declines. Developmental Comparative Immunology, 26(5), 471–479.

Sharma, A., Singla D., Rashid, M. Raghava, G. P. S. (2014). Designing of peptides with desired half-life in intestine-like environment. BMC Bioinformatics, 15, 282.

Sidney, J., Assarsson, E., Moore, C., Ngo, S., Pinilla, C., Sette, A. Peters, B. (2008). Quantitative peptide binding motifs for 19 human and mouse MHC class I molecules derived using positional scanning combinatorial peptide libraries. Immunome Research, 4, 2.

Soltani, S., Keymanesh, K. Sardari, S. (2007). *In vitro* analysis of antifungal peptides: determining the lead template sequence of potent antifungal peptides. Expert Opinion on Drug Discovery, 2(6), 837–847.

Stewart, G. D., Skipworth, R. J. E., Pennington, C. J., Lowrie, A. G., Deans, D. A. C., Edwards, D. R., Habib, F. K., Riddick, A. C. P., Fearon, K. C. H. Ross, J. A. (2008). Variation in dermcidin expression in a range of primary human tumours and in hypoxic/oxidatively stressed human cell lines. British Journal of Cancer, 99(1), 126–132.

Strömstedt, A. A., Pasupuleti, M., Schmidtchen, A. Malmsten, M. (2009). Evaluation of strategies for improving proteolytic resistance of antimicrobial peptides by using variants of EFK17, an internal segment of LL-37. Antimicrobial Agents and Chemotherapy, 53(2), 593–602.

Sturniolo, T., Bono, E., Ding, J., Raddrizzani, L., Tuereci, O., Sahin, U., Braxenthaler, M., Gallazzi, F., Protti, M. P., Sinigaglia, F. Hammer, J. (1999). Generation of tissue-specific and promiscuous HLA ligand databases using DNA microarrays and virtual HLA class II matrices. Nature Biotechnology, 17(6), 555–561.

Thangamani, S., Younis, W. Seleem, M. N. (2015). Repurposing ebselen for treatment of multidrug-resistant staphylococcal infections. Nature Scientific Reports, 5.

Turillazzi, S., Mastrobuoni, G., Dani, F. R., Moneti, G., Pieraccini, G., la Marca, G. Dapporto, L. (2006). Dominulin A and B: two new antibacterial peptides identified on the cuticle and in the venom of the social paper wasp *Polistes dominulus* using MALDI-TOF, MALDI-TOF/TOF, and ESI-ion trap. Journal of the American Society for Mass Spectrometry, 17(3), 376–383.

Wang, G., Li, X., Wang, Z. (2016). APD3: the antimicrobial peptide database as a tool for research and education. Nucleic acids research, 44 (D1), D1087–D1093.

Wang, P., Sidney, J., Dow, C., Mothé, B., Sette, A. Peters, B. (2008). A systematic assessment of MHC class II peptide binding predictions and evaluation of a consensus approach. PLoS Computational Biology, 4(4), e1000048.

Wang, P., Sidney, J., Kim, Y., Sette, A., Lund, O., Nielsen, M. Peters, B. (2010). Peptide binding predictions for HLA DR, DP and DQ molecules. BMC Bioinformatics, 11, 568.

Wardenburg, J. B., Williams, W. A. Missiakas, D. (2006). Host defenses against *Staphylococcus aureus* infection require recognition of bacterial lipoproteins. Proceedings of the National Academy of Sciences, 103(37), 13831–13836.

WHO. (2014). Antimicrobial Resistance: Global Report on Surveillance. France.

Wickham, H. (2009). Ggplot2: Elegant graphics for data analysis. Springer-Verlag, New York.

Yang, X., Hu, Y., Xu, S., Hu, Y., Meng, H., Guo, C. Wang, H. (2013). Identification of multiple antimicrobial peptides from the skin of fine-spined frog, *Hylarana spinulosa* (Ranidae). Biochimie, 95(12), 2429–2436.

Yeaman, M. R. Yount, N. Y. (2003). Mechanisms of antimicrobial peptide action and resistance. Pharmacological reviews, 55(1), 27–55.

Yeung, A. T., Gellatly, S. L. Hancock, R. E. (2011). Multifunctional cationic host defence peptides and their clinical applications. Cellular and Molecular Life Sciences, 68(13), 2161–2176.

Yu, G., Baeder, D. Y., Regoes, R. R. Rolff, J. (2016). Combination effects of antimicrobial peptides. Antimicrobial agents and chemotherapy, 60(3), 1717–1724.

Zhang, L. J. Gallo, R. L. (2016). Antimicrobial peptides. Current Biology, 26(1), R14–R19.

Zhao, H., Zhou, J., Zhang, K., Chu, H., Liu, D., Poon, V. K. M., Chan, C. C. S., Leung, H. C., Fai, N., Lin, Y. P., Zhang, A. J. X., Jin, D. Y., Yuen, K. Y. Zheng, B. J. (2016). A novel peptide with potent and broad-spectrum antiviral activities against multiple respiratory viruses. Scientific reports, 6, 22008.

Zurek, O. W., Pallister, K. B. Voyich, J. M. (2015). *Staphylococcus aureus* inhibits neutrophil-derived IL-8 to promote cell death. The Journal of Infectious Diseases, 212(6), 934–938.

